# Mutational scan inferred binding energetics and structure in intrinsically disordered protein CcdA

**DOI:** 10.1101/2022.04.08.487678

**Authors:** Soumyanetra Chandra, Kavyashree Manjunath, Aparna Asok, Raghavan Varadarajan

## Abstract

Unlike globular proteins, mutational effects on the function of Intrinsically Disordered Proteins (IDPs) are not well-studied. Deep Mutational Scanning of a yeast surface displayed mutant library yields insights into sequence-function relationships in the CcdA IDP. The approach enables facile prediction of interface residues and local structural signatures of the bound conformation. In contrast to previous titration-based approaches which use a number of ligand concentrations, we show that use of a single rationally chosen ligand concentration can provide quantitative estimates of relative binding constants for large numbers of protein variants. This is because the extended interface of IDP ensures that energetic effects of point mutations are spread over a much smaller range than for globular proteins. Our data also provides insights into the much-debated role of helicity and disorder in partner binding of IDPs. Based on this exhaustive mutational sensitivity dataset, a model was developed to predict mutational effects on binding affinity of IDPs that form alpha-helical structures upon binding.

## Introduction

Deciphering sequence function relationships in proteins has been one of the key agendas in biological research in the last few decades. Successful prediction of mutational effects on protein function has important implications for studying genetically driven human diseases as well as in protein engineering and design. The advent of Next Generation sequencing (NGS) and high throughput screening techniques, allowing simultaneous probing of large numbers of protein variants, has greatly facilitated investigations of genotype-phenotype relationships in proteins. Popularly dubbed as Deep Mutational Scanning (DMS) (Fowler and Fields, 2014), this high throughput approach commonly employs screening platforms such as cell based *in vivo* phenotypic assays (Adkar et al., 2012; Hietpas et al., 2011; Jiang et al., 2013; Melamed et al., 2013; Roscoe et al., 2013) or display platforms like yeast surface display (Boder and Wittrup, 1997; Heyne et al., 2020), phage display (Ernst et al., 2010; Ravn et al., 2013), mammalian surface display(Forsyth et al., 2013) and ribosome display (Larman et al., 2012). DMS methods simultaneously evaluate functional features of large libraries, by exploiting variant enrichment following selection, or distribution of the variants amongst different bins based on a functional attribute. Several groups have successfully studied mutational effects on protein-protein interactions (PPIs)(DeBartolo et al., 2012; Ernst et al., 2010; Fowler et al., 2010; Fujino et al., 2012; McLaughlin et al., 2012; Starita et al., 2013; Whitehead et al., 2012) using DMS and attempted epitope mapping(Van Blarcom et al., 2015; Datta et al., 2020; Doolan and Colby, 2015; Kowalsky et al., 2015; Mata-Fink et al., 2013; Najar et al., 2017) as well as quantifying interactions for large numbers of variants(Adams et al., 2016; Jenson et al., 2018).

There has been a plethora of studies employing DMS in globular and structured proteins. In contrast, how amino acid sequence specifies the disorder and yet codes for a biologically relevant function in Intrinsically Disordered Proteins (IDPs) still remains a puzzle. A large part of the genome encodes proteins with functionally important, intrinsically disordered regions (IDRs) (Uversky and Dunker, 2010). IDP(R)s lack well-folded structure in their free, native form. Mutations in the IDRs have been implicated in several human diseases including cancer and neurodegenerative ailments (Uversky, 2014; Uversky et al., 2008). The IDRs are often responsible for binding target proteins with high specificity, and are known to become structured upon binding in several cases (Dyson and Wright, 2005; Mittag et al., 2010). A large number of IDRs fold into helices upon binding, as evident from available structures of IDP-target complexes.

To probe residue specific contributions towards function of an intrinsically disordered protein we studied CcdA, a 72 residue protein antitoxin coded by the *ccdA* component of *ccdAB*, an *E. coli* F-plasmid borne type II toxin-antitoxin system (Miki et al., 1984). A cognate toxin protein, CcdB is expressed from the same operonic system. CcdB exerts its toxicity and mediates cell death by binding to bacterial DNA Gyrase as well as its corresponding DNA adduct (Bernard and Couturier, 1992; Critchlow et al., 1997; Dao-Thi et al., 2005), in the absence of the CcdA antitoxin. The antitoxin CcdA, when present, prevents cell death by binding tightly to CcdB and preventing its binding to Gyrase. CcdA can also rejuvenate Gyrase by extracting CcdB from both the binary CcdB-Gyrase and ternary CcdB-Gyrase-DNA complexes(Aghera et al., 2020; De Jonge et al., 2009; Maki et al., 1996). CcdA has a structured, DNA-binding, N-terminal domain (residues1-39) and an intrinsically disordered C-terminal domain (residues 40-72). The CcdA N-terminal domain is responsible for dimerization (Madl et al., 2006; Van Melderen et al., 1996) and binds to the operator/promoter region of the *ccdAB* operon, playing an important role in autoregulation of *ccdAB* transcription (Afif et al., 2001; Dao-Thi et al., 2002; Vandervelde et al., 2017). The CcdA C-terminal domain, responsible for binding the cognate toxin CcdB, is intrinsically disordered and lacks structure in its free form(Bernard and Couturier, 1991; Dao-Thi et al., 2002; De Jonge et al., 2009; Madl et al., 2006). The available crystal structure of the CcdA C-terminus bound to CcdB dimer reveals that the CcdA C-terminal domain folds into a bent alpha helical region (residues 39-63) and an extended arm(residues 64-72), upon binding to CcdB (De Jonge et al., 2009). The availability of extensive structural information for CcdA in its bound form makes it an ideal model candidate for performing saturation mutagenesis, with the aim of understanding functional consequences of sequence manipulation in an IDP.

We present here a DMS study of CcdA, which explores mutational effects in an IDP on its interaction with a protein partner, using a facile binding energetics quantification methodology, BAMseq (<u>B</u>inding <u>A</u>ffinity <u>M</u>easurement using deep <u>seq</u>uencing). The BAMseq methodology integrates site-directed saturation mutagenesis and deep sequencing with Fluorescence Activated Cell Sorting (FACS) coupled to Yeast Surface Display (YSD). Unlike most other affinity measuring DMS methods, BAMseq uses the ratio of fluorescence signal of ligand binding to that of the surface expression of query protein, as a measure of ligand binding affinity. BAMseq allows parallel high-throughput estimation of binding affinity for diverse and numerous variants of a protein at a single ligand concentration, in contrast to titration based approaches which examine binding at multiple ligand concentrations (Adams et al., 2016). We successfully scored ~1290 single mutants of CcdA based on their relative binding affinity for CcdB. These scores helped elucidate residue specific contributions to CcdA function and identified the CcdA residues crucial for CcdB binding. We also deduced the apparent dissociation constant, K_d_ and change in Gibbs Free Energy of binding, ΔG°_bind_ from the DMS scores for all single-site CcdA mutants in the library. We successfully predicted local secondary structural signatures for the intrinsically disordered CcdA C-terminal domain in its bound form, solely from analysis of the mutational effects across the protein length. This suggests that DMS data can be used to improve or validate structural predictions for IDP bound conformations. This is important, because while there have been revolutionary advancements in deep learning based structural predictions of proteins (Baek et al., 2021; Jumper et al., 2021), IDP structural predictions by these methods have been deemed unreliable (Chakravarty and Porter, 2022; Ruff and Pappu, 2021; Strodel, 2021). Based on the mutational tolerance scores for CcdA single variants, we also developed an interaction weighted mutational penalty-based model, IPApred (Intrinsically disordered <u>P</u>rotein <u>A</u>ffinity <u>pred</u>iction), to predict mutational effects on binding affinity for IDPs involved in PPIs. IPApred showed encouraging results when applied to prediction of mutational effects on partner binding in a number of IDPs.

## Results

### CcdA SSM Library Probed by FACS coupled to YSD

A single-site saturation mutagenesis (SSM) library of all possible single mutants of CcdA, where all 72 positions were individually mutated to all other amino acids, was constructed and cloned in the yeast surface display vector, pETcon (Addgene plasmid # 41522). The CcdA molecules were displayed on the yeast surface as a genetic fusion to Aga-2p, a yeast surface protein (Swers et al., 2004). The surface expression level of CcdA variants was monitored by the epitope tag c-myc fused to its C-terminus. CcdB binding was probed using biotinylated CcdB by FACS (Fig 1).We observed that single mutations in CcdA protein generally fail to completely abolish CcdB binding (Fig 2A-B). The available complex structure of the CcdA C-terminal domain bound to CcdB (PDB ID:3G7Z) (De Jonge et al., 2009) (Fig 2C-D)shows an extensive interface surface area of ~2850Å^2^ between the CcdA C-terminus and the CcdB dimer, explaining the nominal effects of CcdA single mutants on binding energetics.

**Figure 1.**
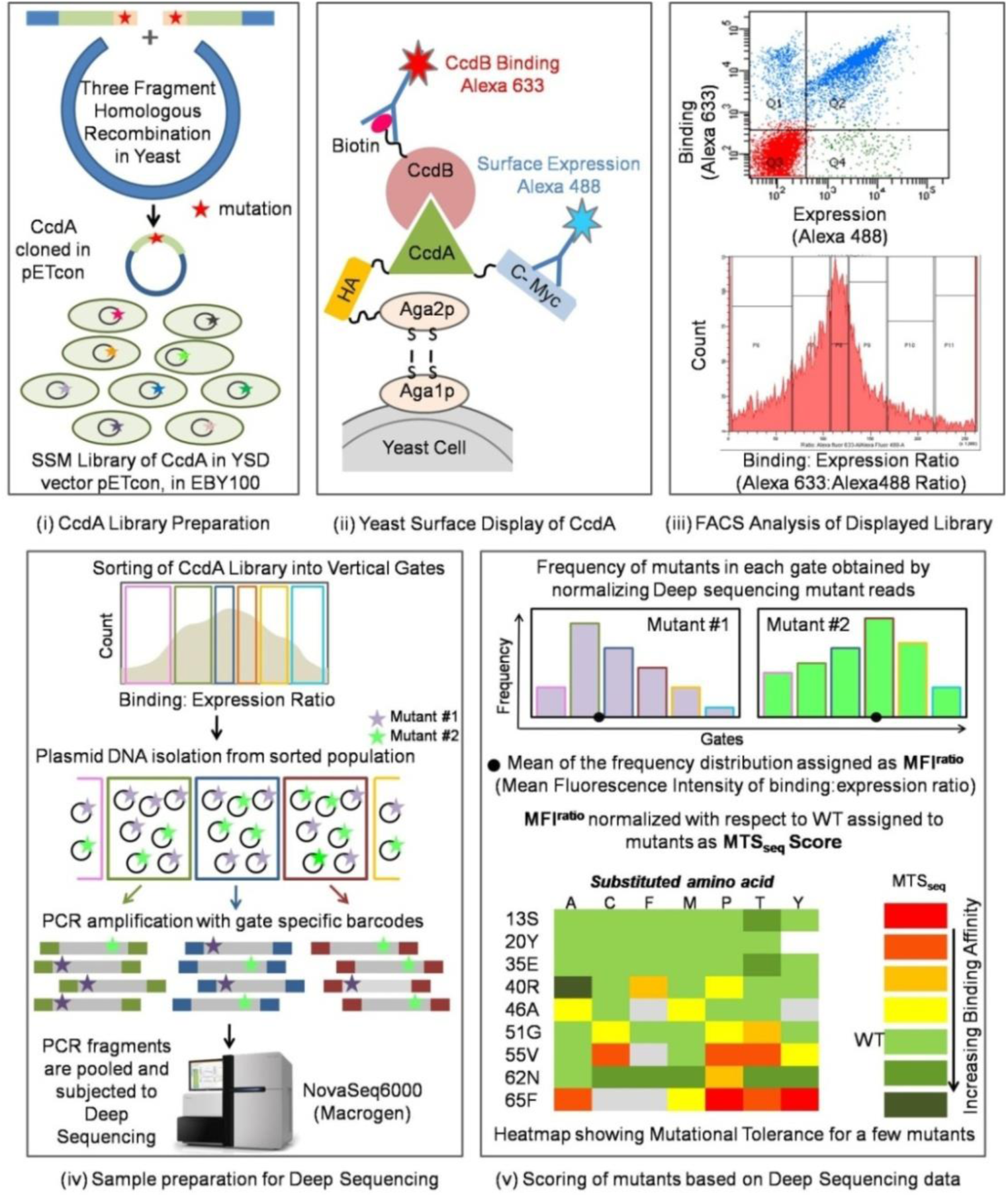
Detailed workflow of deep mutational scanning employed in BAMseq. A single-site saturation mutagenesis library of CcdA in yeast surface display vector pETcon was constructed using three fragment homologous recombination in yeast strain, *S. cerevisiae*. This library was then displayed on the yeast surface as a fusion to the Aga2p surface protein and probed by FACS for its surface expression as well as binding to its target partner, CcdB. The library of CcdA variants is then sorted into vertical bins across an axis representing CcdB binding affinity. After introducing bin specific barcodes, the mutants from different bins are pooled and subjected to deep sequencing. After normalisation of the obtained raw data representing the mutant distribution across the affinity bins, a mutational tolerance score (MTS_seq_) is assigned to each mutant (see Methods), which allows both ranking of mutants based on affinity as well as undestanding residue specific contributions to the binding enegetics.

**Figure 2.**
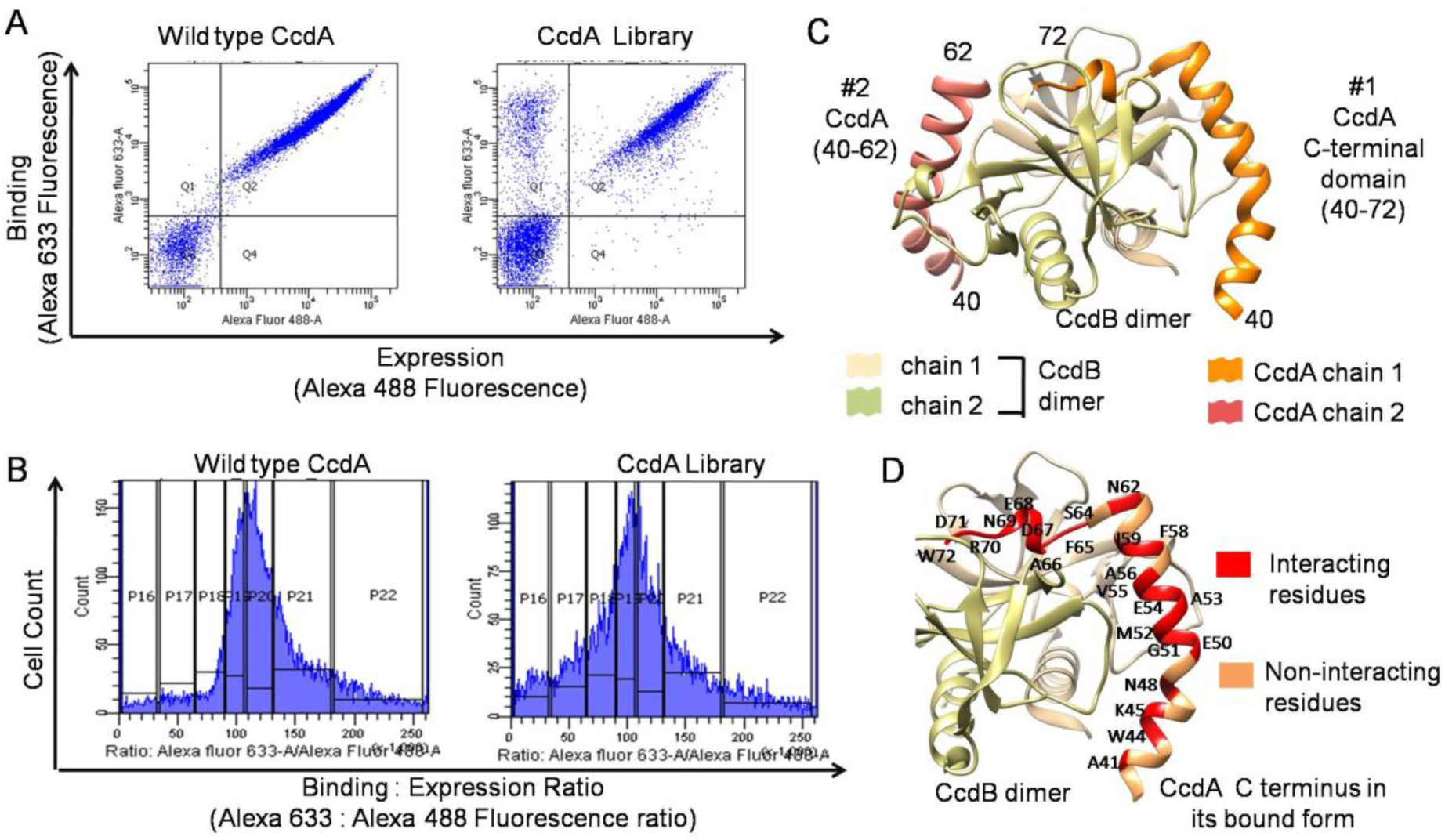
CcdB binding activity of WT and mutant CcdA. (A) FACS analysis of the surface displayed CcdA WT (left) and SSM Library (right) showing CcdB binding versus surface expression plots. The surface expression and CcdB binding of CcdA are monitored by Alexa-488 and Alexa-633 fluorescence intensity respectively. Most CcdA variants have similar binding affinity to WT as evident from the similarity of the two plots. (B) Cell count as a function of CcdB Binding : Expression Fluorescence Intensity ratio for CcdA Wild type and SSM library. This allows for sorting based on relative affinity towards CcdB. The CcdA library was sorted into seven vertical bins, P16-P22. (C) The crystal structure of the intrinsically disordered C-terminal domain of CcdA in complex with the CcdB dimer (PDB ID : 3G7Z). The CcdB dimer has two asymmetric binding sites for CcdA with differing affinity, one binding the full 40-72 residue C-terminal domain with high affinity (CcdA chain 1) and the other binding to only the 40-62 stretch of the C-terminal domain of a second CcdA molecule with lower affinity (CcdA chain 1). (D) A closer view of the high affinity binding site between the CcdA C-terminal domain and the CcdB dimer, with CcdB interacting and non-interacting residues, as inferred from the crystal structure, highlighted in red and peach respectively. The interacting residues are labeled (WT residue identity and position).

### Scoring Mutants for their Binding Affinity and Surface Expression using FACS coupled to deep sequencing

One way to quantify affinity is to monitor the diagonal axis of the Binding versus Expression FACS plot, sorting the library into parallel diagonal gates, as used in SORTCERY(Reich et al., 2015, 2016). Affinity quantification from the diagonal axis is both experimentally and mathematically intricate and is unsuitable for libraries where mutational effects are quite moderate, as seen in the case of CcdA (Fig 2A). Previously, binding affinity for a variant library was also quantified by sorting based on the binding fluorescence, at multiple target concentrations in a titration like strategy(Adams et al., 2016). The requirement of sorting into multiple gates at multiple target concentrations makes this method both labour intensive and expensive. Instead we used a simpler approach, measuring the cell count as a function of Binding:Expression ratio (Alexa 633:Alexa 488 fluorescence) at a single ligand concentration, where the x-axis (Binding:Expression ratio) can be taken as an approximate measure of the CcdB binding affinity of displayed CcdA molecules (Fig 2B). Sorting the population into vertical bins across this x-axis separates the library into compartments with varying degrees of binding affinity. After introduction of bin specific barcodes that provide bin identification, the *ccdA* amplicon DNA from all the sorted populations was pooled and subjected to deep sequencing (MacrogenNovaSeq6000). The raw read distribution across bins in the deep sequencing data was used to reconstruct the frequency distribution of each mutant across the vertical bins. We verified for a small number of mutants, that the reconstituted frequency distribution obtained from deep sequencing data analysis resembles the individual FACS profiles of the respective single mutants (Supplementary Fig S1A-B).

We recently used a similar approach to estimate mean fluorescence intensity, MFI of CcdB mutants using YSD and showed that this simplified approach yields MFI values similar to more complicated Maximum Entropy based approaches (Ahmed et al., 2022).The MFI value of the frequency distribution across the Binding: Expression ratio axis is assigned as MFI^ratio^ for ~1290 single mutants of CcdA (94 % of the total library diversity) (see Methods and Fig 1). The deep sequencing and MFI^ratio^ assignment were carried out in two biological replicates, which were highly correlated (Supplementary FigS1C). The MFI^ratio^ values were normalized with respect to the WT MFI^ratio^ and averaged across the replicates to obtain the deep sequencing derived Mutational Tolerance Scores (MTS_seq_) for CcdA mutants. The MTS_seq_ score provides a measure of the CcdB relative binding affinity.

All mutations bearing synonymous codons with respect to the WT sequence were neutral in the YSD binding assay. We also found that mutants coded by degenerate codons had similar inferred relative binding affinity, indicating that the mutational effects on binding observed in surface displayed CcdA are solely due to the primary amino acid sequence and not the nucleic acid sequences. We also characterized surface expression of the CcdA library by sorting into bins, based on the fluorescence associated with the C-terminal c-myc tag (Alexa 488 fluorescence) (Supplementary Fig S1D). Expression scores obtained from deep sequencing for the CcdA mutants showed absence of significant mutational effects on surface expression (Supplementary Fig S2), unlike effects seen in case of a notable fraction of yeast surface displayed globular protein variants(Ahmed et al., 2021; Park et al., 2006). This suggests that sequence variation has a minimal effect on IDP proteolysis, which plays an essential role in TA system activation under stress conditions. Unlike globular proteins, where mutations affecting protein structure and stability may lead to increased aggregation and/or proteolysis and therefore lower expression levels on the yeast surface, expression profiles for intrinsically disordered proteins appear to be minimally affected by mutations. Surprisingly, certain mutants at C-terminal domain positions (especially residues 55-59) show an elevated expression level on the yeast surface (Supplementary Fig S2), indicating that the WT sequence of this C-terminal domain stretch might be responsible for promoting disorder but is otherwise destabilizing, and is rescued by mutations.

### Experimental validation of BAMseq methodology

We experimentally determined the dissociation constant, K_d_for twenty individual single mutants and WT CcdA (Supplementary Figs S3A and S4) by CcdB titration of surface displayed CcdA molecules, using FACS. Surface displayed WT CcdA exhibited a K_d_ value of 0.25±0.05 nM for CcdB binding (Supplementary Fig S3A and Supplementary Table S1), in our YSD system. It has been previously reported that the CcdB dimer has two non-identical sites that bind two molecules of CcdA with distinct binding affinities, one binding the full length C-terminal domain of CcdA with a K_d_ of 20 pM, while another binds only residues 40-61 of another CcdA C terminal domain with a K_d_ of 13 μM (De Jonge et al., 2009).However, recent Surface Plasmon Resonance (SPR) and fluorescence kinetic studies of CcdA-CcdB_2_ interactions showed a K_d_ value of ~0.3 nM for the high affinity site (Aghera et al., 2020) consistent with the present value. Since full length CcdA protein is intrinsically disordered, it is unstable and aggregation prone upon purification (Dao-Thi et al., 2002), which significantly complicates investigation of its interaction with partners using classical techniques like ITC (Isothermal Titration Calorimetry) or SPR. Instead, the YSD based titration experiment used here serves as a facile approach to experimentally quantify CcdB binding affinity for a finite number of individual CcdA mutants. Several studies have previously established that binding constant values derived from YSD based FACS titration agree well with those obtained from classical methods like ITC, SPR and BLI as well as other techniques such as equilibrium competition titrations, fluorescence quenching titrations and stopped flow kinetics consistently(Bacon et al., 2020; Lipovsek et al., 2007; Rathore et al., 2018; Uchański et al., 2019).

The change in Gibbs free energy of CcdB binding (ΔG°_bind_) was calculated for each of the twenty CcdA mutants using the fitted K_d_ values (Supplementary Table S1). These experimentally determined ΔG°_bind_ values correlate well with the DMS derived MTS_seq_ scores (Pearson correlation, r= −0.84) (Supplementary Fig S3B) confirming the accuracy and precision of mutational scores derived from our BAMseq methodology. The experimental dissociation constant (K_d_) values show a non-linear relationship to the MTS_seq_ scores and fit better to an inverse first order equation (Supplementary Fig S3C), consistent with the ratio of binding to expression on yeast cell surface being inversely proportional to the K_d_ values. Theoretically, binding: expression fluorescence signal is found to be proportional to [C] / (K_d_ + [C] + [A])(see Supplementary Methods), where K_d_ = dissociation constant, [C] = CcdB (ligand) concentration used in the FACS experiment and [A]= concentration of free CcdA. Simulation of theoretical curves helps to determine the ideal ligand concentrations that can be used in BAMseq experiments for estimating binding affinities accurately (Supplementary Fig S5).

### Determination of apparent dissociation constant (K_d_) for all single-site mutants of CcdA

We further developed the BAMseq method to allow determination of binding constants from the DMS derived MTS_seq_ scores, since the MTS_seq_score only provides a relative measure of the CcdB binding affinities of CcdA mutants. The highly correlated MTS_seq_ scores and experimental ΔG°_bind_ values of twenty experimentally validated mutants provide the necessary standard curve(Supplementary Fig S3B), facilitating extrapolation of apparent ΔG°_bind_ and apparent K_d_ for mutants from the MTS_seq_ scores. A similar method of extrapolation of binding energetics from DMS has been described previously(Jenson et al., 2018). Using this method, we could assign apparent ΔG°_bind_ and K_d_ values for all ~1290 CcdA single mutants that lack prior experimental binding energetic information. The apparent ΔG°_bind_^seq^ predicted by BAMseq show a normal distribution with a shoulder at the lower CcdB binding affinity range, indicating a distinct population of binding defective CcdA mutants (Supplementary Fig S3D).

### Mutational sensitivity landscape for CcdA confirms the redundancy of the N-terminal domain in CcdB binding activity

In our CcdA DMS study, the N-terminal region (1-39) shows high tolerance to mutations, with MTS_seq_ values close to that of WT CcdA, except for a few mutants that display modest impairment in CcdB binding (Fig 3A). Minor binding defects upon mutation occur primarily at buried positions (Fig 3A), when mapped onto the available DNA bound, CcdA dimer structure (Madl et al., 2006). This may result from disruption of the hydrophobic core upon mutations in the structured N-terminal domain, leading to the possible aggregation of the displayed full length CcdA protein on the yeast surface, consequently decreasing CcdB binding. It is noteworthy that these mutations show no distinguishable effects on surface expression (Supplementary Fig S2) indicating an absence of enhanced proteolysis or degradation of the variant protein on the yeast surface. Our DMS results clearly indicate that mutations in the N-terminal residues fail to impair the CcdB binding activity and therefore can be inferred to have minimal contributions to the binding energetics or binding interface, consistent with the NMR and proteolysis studies of CcdB bound CcdA molecules, carried out previously (Madl et al., 2006). We also see no effects on CcdB binding of the displayed CcdA molecule upon mutations at positions involved in dimerization of the N-terminal domain (Madl et al., 2006; Van Melderen et al., 1996) (Fig 3A), confirming that dimerization of the N-terminus of CcdA is non-essential for CcdB binding.

**Figure 3.**
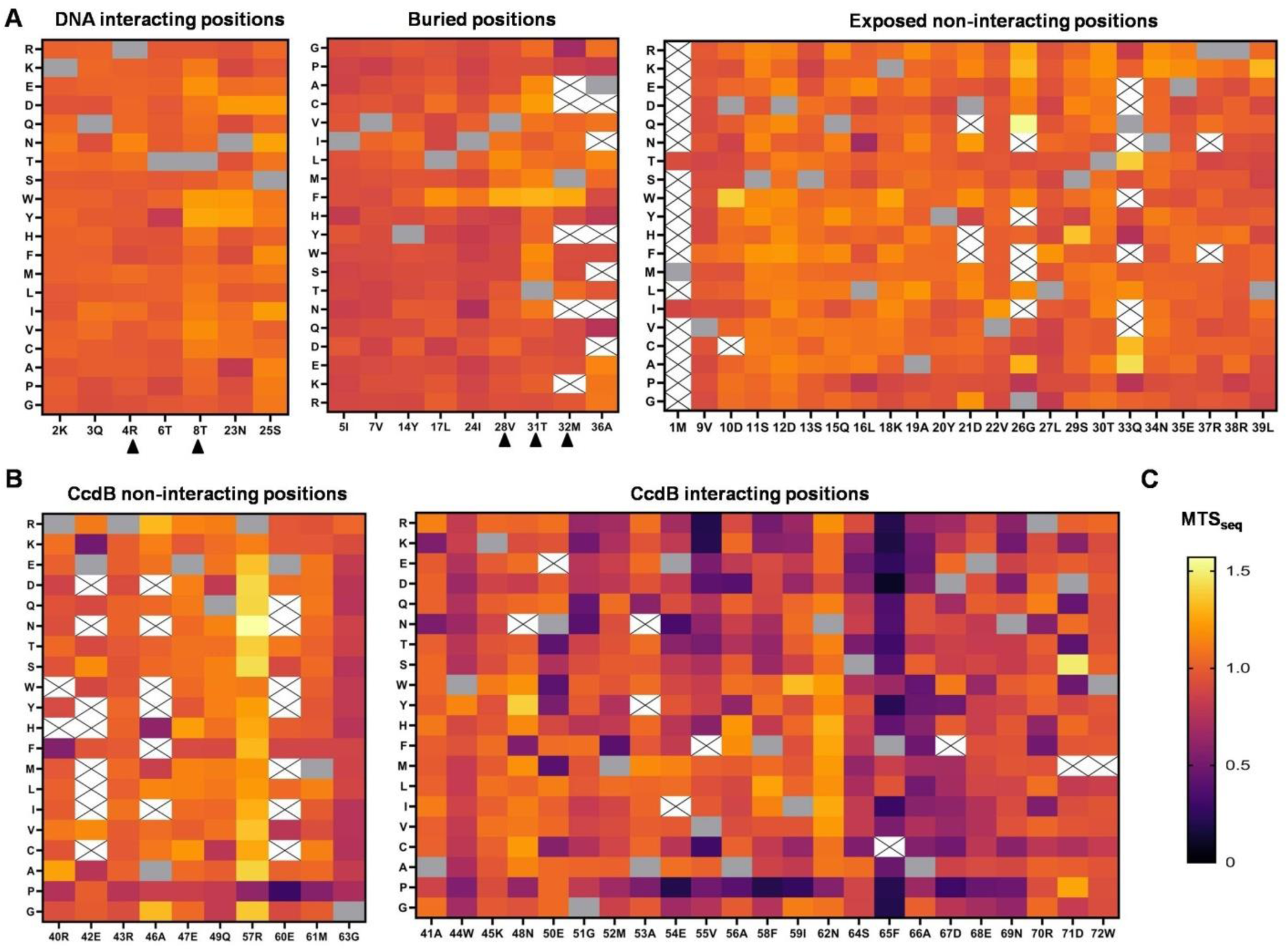
Mutational tolerance of CcdA single mutants. Mutational tolerance scores for (A) the structured N-terminal DNA binding domain of CcdA and (B) the intrinsically disordered, CcdB binding C-terminal domain of CcdA. The N-terminal domain residues are classified as DNA interacting, buried and non-interacting sites based on the DNA bound dimeric CcdA N-terminal domain structure (PDB ID :2H3C)(Madl et al., 2006). The positions known to be involved in the N-terminal domain dimerization in the DNA bound CcdA dimer structure, are marked with black triangles. Residues in the C-terminal disordered domain of CcdA are classified as CcdB interacting and non-interacting based on the CcdAB complex structure (PDB ID :3G7Z). (C) The color key used depicts the ranges of MTS_seq_ scores representing CcdB affinity. WT CcdA has MTS_seq_ score of 1. The poor binders with lower tolerance scores are shown in dark colors, while lighter colors depict marginally better binders. Grey cells and blank cells (marked with X) indicate WT residue and mutants with no available data, respectively.

The detrimental mutational effects on binding activity, as revealed by our DMS study of the CcdA, are restricted largely to the C-terminal intrinsically disordered domain (residues 40-72) and generally to the residues inferred to be CcdB interacting, from the available CcdA-CcdB complex structure(De Jonge et al., 2009) (Fig 3B). CcdA typically exhibits a high mutational tolerance throughout its length, barring a few positions, which can therefore be deduced as functionally important, CcdB binding residues.

### Prediction of binding interface from mutational landscape of an IDP

To establish DMS as a viable method for prediction of interface residues in IDPs, we studied the relationship between overall mutational tolerance of CcdA C-terminal residues, with several structural parameters such as the change in Accessible Surface Area upon binding (ΔASA), residue depth (both commonly used to quantify residue specific contribution to interaction) or the number of bonds and interactions derived from the available complex structure of CcdA-CcdB (Supplementary Table S2). ΔASA is calculated by subtracting the accessible surface area (Lee and Richards, 1971) of CcdA in complex with CcdB from the accessible surface area of free structured CcdA. The Residue Depth is a measure of the depth of a residue from the exposed surface (Chakravarty and Varadarajan, 1999; Tan et al., 2011), with a higher depth in the bound form signifying interaction with partner. The mutational tolerance averaged across all substitutions (MTS_seq_^avg^) of each residue position correlates fairly well with the respective ΔASA, number of interactions and residue depth (Fig 4A-C).

**Figure 4.**
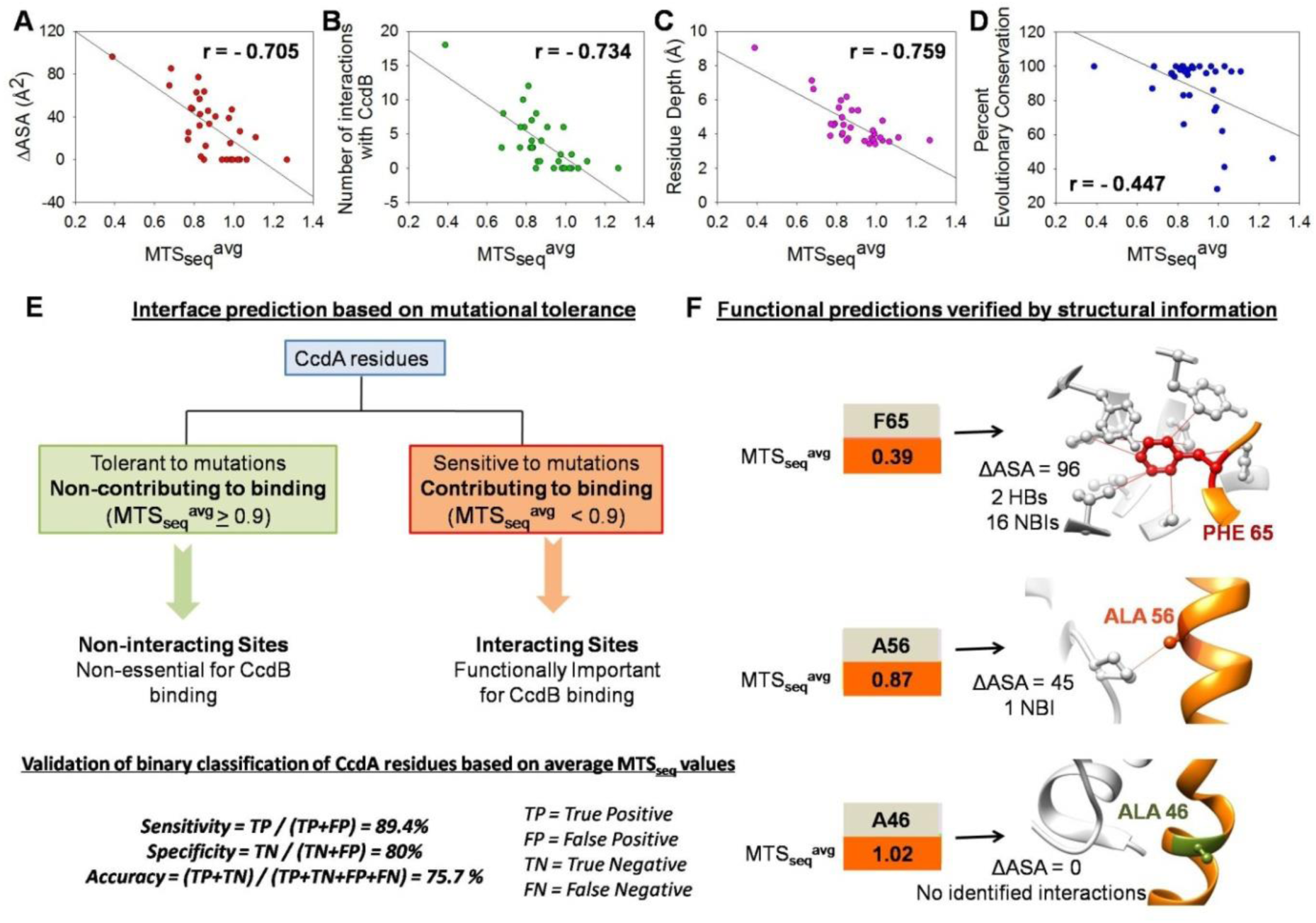
Prediction of interface residues from Deep Mutational Scanning of CcdA. Correlations of the average mutational tolerance scores (MTS_seq_^avg^) for residues in the CcdA C-terminal domain with (A) ΔASA (B) number of interactions each position is involved in with CcdB (C) Residue Depth respectively, calculated using the CcdAB complex structure (PDB ID: 3G7Z) and (D) the percent evolutionary conservation at each position across bacterial species. r is the Pearson Correlation Coefficient. The MTS_seq_^avg^ scores were calculated by averaging the MTS_seq_ for all substitutions at each position and depict the average effect of mutations at each position. (E) Flowchart for prediction of functionally important residues in CcdA from the positional MTS ^avg^ scores (F) Validation of interface predictions by structural analysis. HB and NBI refers to number of H-bonds and non-bonded interactions respectively that the WT residue forms with CcdB (grey) in the crystal structure of the complex.

Functionally important residues in proteins are generally found to be conserved across species in order to maintain optimal activity. Surprisingly, the MTS_seq_^avg^ of CcdA C-terminal residues show only a modest correlation (r = −0.45) with the percent conservation of the *E. coli* WT residues in CcdA molecules across prokaryotes (Fig 4D, Supplementary Table S2). At some positions, mutations that are poorly tolerated in *E. coli* CcdA are found in other species. This might be due to co-evolution, with cognate CcdB harboring compensating mutations. In case of *E.coli* CcdB protein, CcdA interacting residues are also found to be poorly conserved across homologs.

Statistical k-means clustering of average mutational effects in CcdA into two classes, yields class-1 (similar to WT) and class-2 (lower binding affinity) having mean MTS_seq_ values of 1.02 ± 0.152 and 0.49 ± 0.165 respectively. We categorize mutants having MTS_seq_ lower than 0.9 (≈ mean^class2^+ 2.5 σ^class2^) as CcdB binding defective. A high cutoff value for this classification was chosen deliberately to allow identification of mutants with moderate binding defects, as mutational effects are observed to be typically mild in case of the CcdA library. CcdA residues can also be classified as CcdB interacting and non-interacting based on the ΔASA values obtained from the CcdA-CcdB structure (De Jonge et al., 2009). When compared to these structure based classes, the contributing and non-contributing classes of residues in CcdA identified solely based on a MTS_seq_ ^avg^ cutoff of 0.9 yields reasonable sensitivity (89%), specificity (80%) and accuracy (76%) calculated as described previously(Bhasin and Varadarajan, 2021; Tripathi et al., 2016) (Fig 4E-F).

While all the structure inferred non-interacting positions with the exception of Gly 63 (discussed later) were mutation tolerant, only 75% of the structure inferred CcdB interacting positions showed a significant decrease in CcdB binding activity upon mutation (Supplementary Fig S6).Despite the high residue burial upon binding and presence of contacts observed in the complex structure, these residues in CcdA exhibit high tolerance to mutations and thus minimal contribution to CcdB binding. CcdA mutational tolerance is non-uniform along its length. With the exception of W44, the 40-49 residue and the 60-64 residue stretches, as well as the C-terminal 71D and 72W residues in the CcdA C-terminal domain show high tolerance to mutations, irrespective of individual residue specific interactions with CcdB identified from the crystal structure (Supplementary Fig S6). This indicates that the residue contribution to CcdB binding in CcdA is higher at the interior of the interface and particularly at regions containing adjacently placed functionally important sites. Our data therefore suggests that in case of extensive interfaces as seen in CcdA, prediction of mutational effects solely from residue specific interactions derived from the crystal structure is likely to be inaccurate.

### Dissecting the plausible role of intrinsic disorder and helicity in CcdB binding of CcdA

The CcdA C-terminal domain is intrinsically disordered and folds into a helical structure upon CcdB binding. We found that mutational tolerance in CcdA correlates negatively with predicted disorder (predicted using IUPred algorithm (Mészáros et al., 2018)) especially for mutations at CcdB interacting sites of CcdA (Supplementary Figure S7A-B). Intrinsic disorder might be essential for CcdA to maintain a disordered state, required for fast and efficient antitoxin degradation necessary in TA modules, and also to allow binding to multiple partners as observed in case of many IDPs. However, the present data indicates that increased disorder leads to impairment in binding affinity towards the partner as well as reduced activity. Similar findings have also been reported in previous DMS studies on IDPs, namely α–synuclein and a transcription factor, Gcn4 (Newberry et al., 2020; Staller et al., 2018). Since many IDPs form helical structure upon partner binding, we studied the relationship between helix propensity and the observed effects of mutations. We observed that amino acid residues that promote helix formation are enriched in the known α-helical stretches in both the N- and C-terminal domains of CcdA (Supplementary Fig S7C). However, we found no apparent relationship between mutational effects on CcdB binding activity in CcdA variants and the relative helix propensity of the mutant amino acid residue with respect to that of the WT residue (Supplementary Fig S7D). If helix formation is a prerequisite for partner binding in IDPs, mutations that decrease helical propensity in these regions can be expected to hinder interactions with partner protein. However, in the case of CcdA the average mutational tolerance of each residue position in CcdA shows only a weak correlation with the helix propensity of the wildtype residues (Supplementary Fig S7E), indicating that partner binding and activity of CcdA do not depend strongly on inherent helix forming capability of the disordered regions.

### Interface Identification by Cysteine and Charged Scanning Methods

A popular method of identifying residue contributions to partner binding in protein interfaces, is Alanine Scanning, where each residue is mutated to alanine to investigate the effect of mutations on protein-protein interactions (PPIs)(Cunningham and Wells, 1989). However Ala scanning performs poorly in many cases(Gray et al., 2017).Ser, Asp, Cys, Asn, His Scanning methodologies have been advocated as alternatives in identification of active site residues in globular proteins or in mapping epitopes (Das et al., 2020; Gray et al., 2017; Kanaya et al., 1990; Pál et al., 2005). We correlated the MTS_seq_ patterns for each of the 20 substituted amino acids with the average residue MTS_seq_ in CcdA. We found charged and cysteine mutants best recapitulated the overall mutational tolerance of the different positions in the CcdA C-terminal domain (Supplementary Fig 8A). Further, the single mutant MTS_seq_ scores for Cys and charged mutants correlate well with the residue depth of CcdA residues in the complex (Supplementary Fig 8B). While the charged substitutions possibly alter the primary interactions at functionally important residues, Cysteine, an aliphatic residue is not expected to alter partner binding. The disruptive effect of Cysteine substitutions predominantly at the interacting positions, might be due to its potential to form intermolecular disulfide bonds, in the oxidizing extracellular environment, masking the interacting face of the extended helical CcdB binding C-terminal domain of CcdA. In contrast, the Ala mutants produce relatively mild effects on binding affinity and fail to distinguish interacting from non-interacting residues in CcdA (Supplementary Fig S8).

### Secondary structural footprint deduced from mutational tolerance pattern

The mutational sensitivity pattern of the CcdA C-terminus derived from BAMseq data shows an apparent oscillating pattern which is clearer for substitutions to charged residues, MTS_seq_ (charged) (Fig 5A-B). It has been previously shown that mutational effects of charged residues on aggregation propensity of the membrane binding amphipathic helix α-synuclein resemble a waveform(Newberry et al., 2020). Though the intrinsically disordered CcdA C-terminal domain is not predicted to be amphipathic based on the hydrophobic moment calculated as described previously (Eisenberg, 1984) or according to estimations by the AMPHIPASEEK web server (Combet et al., 2000), we observe that the effects of charged substitutions on the binding activity in the CcdB interacting and non-interacting face of the CcdA C-terminal helix are clearly distinguishable. We therefore attempted to determine the periodicity in the mutational pattern and thus infer local structural features attained by the intrinsically disordered CcdA upon binding. To correct for the unequal contribution to binding across the length of the CcdA protein, we subtracted the mutational tolerance averaged for charged substitutions, MTS_seq_ (charged) at each position i by the same quantity averaged over a five residue sliding window centered around position i. We tested three, five, seven, and nine residue window averages and chose to use a five-residue window average, since it allows for clear detection of possible phase changes in the waveform. Identification of such discontinuity is essential in correctly predicting structures of IDPs in their partner bound form, where in many cases, the IDPs wrap around the globular partner protein upon binding, and there are drastic angular changes in the protein backbone.

**Figure 5.**
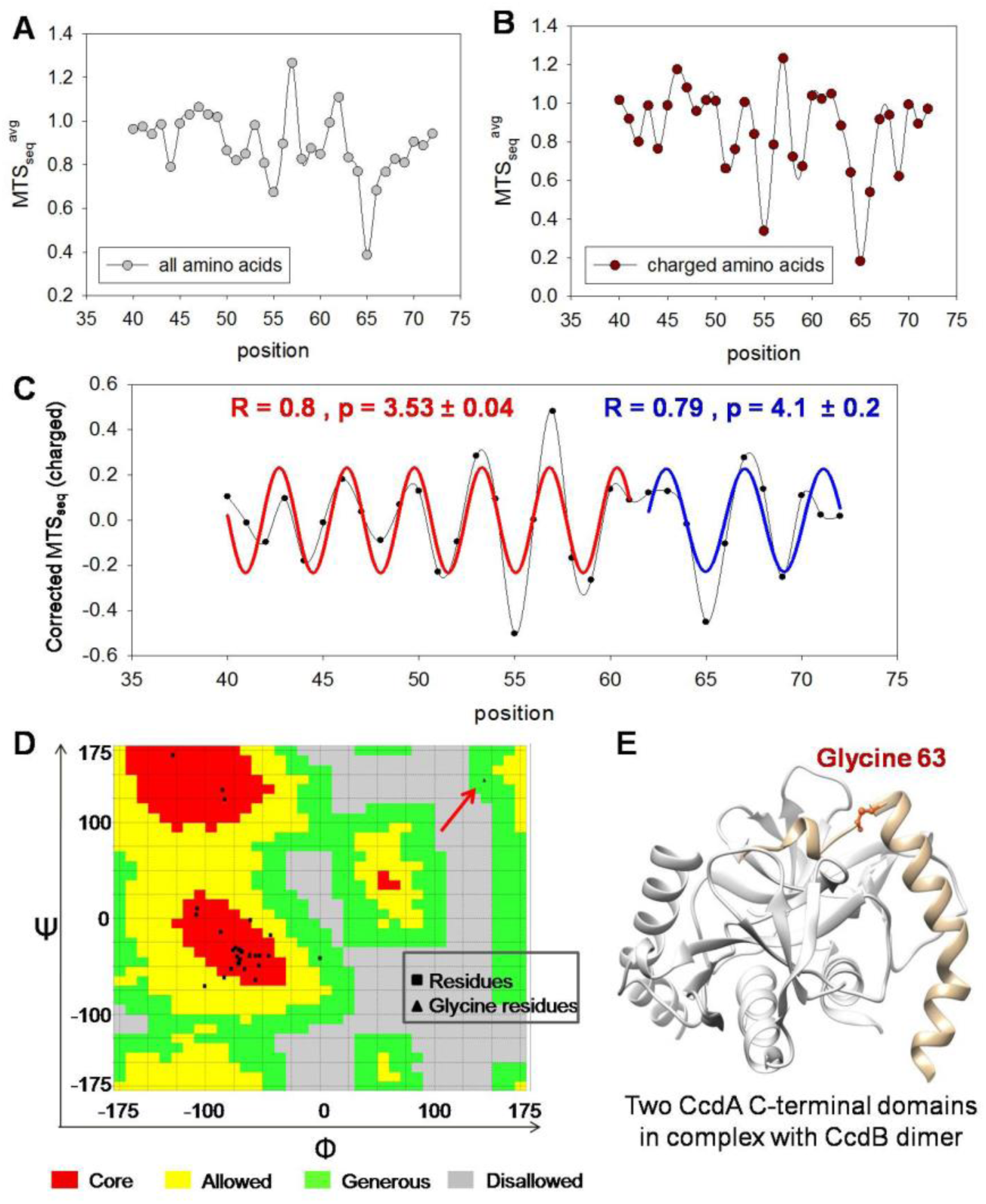
Local structural insights from CcdA C-terminal mutational data. (A) The mutational tolerance score averaged for all substitutions (MTS_seq_^avg^) as a function of residue position in the CcdA C-terminal domain (plotted as a spline curve) displays an oscillating pattern. (B) The mutational tolerance score averaged for charged substitutions, MTS_seq_(charged) as a function of residue position. (C) Corrected mutational tolerance for charged substitutions plotted against position across the length of the CcdA C-terminal domain. The y axis is the difference between averaged mutational tolerance for charged substitutions for each position and the mutational tolerance for charged substitutions averaged over a five residue sliding window [i.e. corrected MTS_seq_(charged) = MTS_seq_(charged) – average MTS_seq_(charged) for a sliding window of five residues], to correct for the uneven distribution of mutational effects across the length of the C-terminal domain. The data shows periodicity and is fitted to a single sinusoidal wave form, y=a sin (2πx/b+c), where a = amplitude, b = periodicity and c = phase. A possible phase change is apparent at residue 63. The two C-terminal domain stretches separated by residue 63, when fitted to two separate wave equations (fitted curves shown in red and blue), show distinct periodicities of 3.54 and 4.1 in the 40-62 region and the 63-72 region respectively. (D) Ramachandran map for all C-terminal domain residues. The arrow shows the G63 torsional angle values highlighted with an arrow. (E) Back bone conformation of Glycine 63 mapped on the structure of the C-terminal domain of CcdA bound to CcdB dimer (PDB ID: 3G7Z).

When fitted to a single sinusoidal wave equation, the corrected MTS_seq_ (charged) shows a poor fit. Upon identification of a possible phase change in the oscillating pattern at residue 63, we fit the corrected charged mutational tolerance values of the two residue segments to separate sinusoidal wave equations. The first 23 residue stretch of the C-terminal domain (residues 40-62) showed a reasonably good fit (R= 0.8) with a periodicity of 3.53± 0.04, consistent with an α-helix containing 3.6 amino acids per turn (Fig 5C). However, the later 10 residue stretch (residues 63-72) fit better to a different waveform with periodicity of 4.1± 0.2, with a good fit (R = 0.79), and a phase difference with the first waveform (Fig 5C). This indicates that the first 23 amino acids (region 40-62) form an α-helix while the next stretch does not, which is consistent with the available structure of CcdB bound CcdA C-terminal domain. Residues 40-62 form a bent helix, while residues 63-72 remain mostly unstructured with a central helical turn at residues 66-69 in the CcdB bound, crystal structure (De Jonge et al., 2009). The troughs in the waveform consist of residue positions 41, 42, 44, 45, 48, 51, 52, 55, 58, 59, 64, 65, 66 and 69 in CcdA, all of which (with the exception of 42) are known to interact with CcdB based on the available CcdA-CcdB crystal structure (De Jonge et al., 2009). This data can be used to further validate or as an alternative approach to examining average mutational sensitivity as described in Figure 4. Only one face of the helix formed upon binding interacts with partner in IDP, resulting in a clear periodicity in the mutational effects. This periodicity when captured in terms of mutational scores allows facile identification of secondary structure and residues that constitute the binding interface. This demonstrates the potential of DMS for elucidation of local structural features of intrinsically disordered proteins and extended interfaces. The amplitude of the mutational tolerance waveform varies across the protein length, owing to differential contributions of individual residues to binding. The MTS_seq_ scores for individual charged substitutions also show a similar oscillating pattern to the averaged values, (Supplementary Fig S9) and can be used to predict periodicity and structure.

### Optimal CcdB binding requires a bent alpha-helix

While all other non-interacting residues are tolerant to mutations, the non-interacting residue Glycine 63 shows a binding defect upon most substitutions (Fig 3B). This is because Glycine 63 has a non-canonical backbone conformation in the bound state (Φ,Ψ = 143, 144) (Fig 5D), which is uncharacteristic of an α-helix, and unachievable by all other amino acid residues. G63 is responsible for breaking the helix (40-62) and allowing the subsequent unstructured tail (64-72) to bend and bind CcdB optimally (Fig 5E). The CcdB dimer is known to have two asymmetrical binding sites for CcdA, one where only the CcdA C-terminal stretch 40-63 binds with low affinity (K_d_ = 13 μM) and the second, where the whole C-terminal 40-72 stretch binds with a high affinity (K_d_ = 20 pM) (De Jonge et al., 2009). The 40-63 stretch as well as the 62-72 stretch have both been shown individually to bind the CcdB dimer with low affinity using ITC experiments(De Jonge et al., 2009). The Glycine residue is thus evidently responsible for providing a flexible hinge between the alpha helical region (39-63) and the extended arm (64-72), a structural feature which appears to be crucial for high affinity CcdB binding. Upon analysis, we further identified similar glycine residues with positive Φ values of torsion angle in antitoxins of many type II TA systems (Table 1), that are positioned specifically to create a drastic angular change in the protein backbone molding the geometry of the typically extended interfaces involved in IDP antitoxins interacting with their cognate toxin partners.

**Table 1.** Predominance of glycine residues with positive torsion angles in bacterial type II antitoxins.

| | Residue/<br>Position<sup>a</sup> | Antitoxin | Complex<br>PDB ID | Chain | TA module | Organism | $\Phi$<br>torsion<br>angle<br>(°)<sup>b</sup> |
| --- | --- | --- | --- | --- | --- | --- | --- |
| 1 | Gly 25 | Maz E | 6A6X | C | MazEF mt9 | *Mycobacterium tuberculosis* | +71 |
| 2 | **Gly 76** | Maz E | 7D2P | E | MazEF | *Deinococcus radiodurans* | +47 |
| 3 | **Gly 138** | Maz E | 6L2A | F | MazEF 4 | *Mycobacterium tuberculosis* | +133 |
| 4 | Gly 35 | Vap B | 6SD6 | A | VapBC | *Shigella sonnei* | +68 |
| 5 | **Gly 44** | Vap B | 6SD6 | A | VapBC | *Shigella sonnei* | +68 |
| 6 | **Gly 67** | Vap B | 3DBO | A | VapBC 5 | *Mycobacterium tuberculosis* | +88 |
| 7 | Gly 9 | Vap B | 6NKL | C | VapBC 5 | *Haemophilus influenzae* | +65 |
| 8 | Gly 35 | Vap B | 6NKL | C | VapBC 5 | *Haemophilus influenzae* | +91 |
| 9 | **Gly 55** | Vap B | 6NKL | C | VapBC 5 | *Haemophilus influenzae* | +117 |
| 10 | Gly 35 | Vap B | 3TND | B | VapBC | *Shigella flexneri* | +68 |
| 11 | **Gly 44** | Vap B | 3TND | B | VapBC | *Shigella flexneri* | +99 |

<sup>a</sup> Glycine residues in the C-terminal toxin binding domain of the antitoxins are highlighted in bold.
<sup>b</sup> $\Phi$ refers to the C-N-C $\alpha$ -C backbone torsion angle for the Glycine residue calculated from complex structures using PDB Goodies (Hussain et al., 2002).

### Performance of available computational predictors of binding affinity changes in mutants

Several commonly used predictors of mutational effect on binding affinity, including BindProfX, PRODIGY, SAAMBE-3D etc fail to make predictions in cases like CcdA (interacting with a dimeric CcdB), where proteins with more than one protomer are involved in the interaction. The predictors mCSM-PPI (Rodrigues et al., 2019), BeAtMuSiC (Dehouck et al., 2013) and MutaBind (Li et al., 2016) do not have this limitation and showed modest correlations with Pearson correlation co-efficient (r) values of 0.49, 0.58 and 0.65 respectively with the experimental ΔΔG°_bind_ (= ΔG°_bind_^mut^-ΔΔG°_bind_^WT^) values for twenty single CcdA mutants (Supplementary Fig S10A-C). However, these predicted mutational effects are significantly larger than observed experimentally, for CcdA. The BAMseq derived apparent ΔG°_bind_^seq^ deduced for all single mutants by BAMseq was compared to predictions by the available computational tool BeAtMuSiC, that predicts binding affinity changes upon all possible single mutations (Dehouck et al., 2013) and a modest correlation with r = 0.45 was observed (Supplementary Fig S10D). Comparison of apparent ΔΔG°_bind_^seq^ deduced for all single CcdA mutants by BAMseq to the predicted ΔΔG°_bind_ values using BeAtMuSiC indicates that many positions with high mutational tolerance are erroneously predicted to have severely depleted binding affinity by BeAtMuSiC, and also the program fails to successfully predict effects in the most mutation sensitive positions (residues 64-67) of CcdA (Supplementary Fig S11). These observations highlight the drawbacks of using the currently available predictors of mutational effects on energetics of PPIs in case of IDPs.

### Prediction of Mutant activity in IDPs based on DMS data

In the present study, we have endeavored to delineate the contribution of every amino residue in the CcdA molecule to its CcdB binding activity. Based on the observed dependence ofmutational tolerance in CcdA on residue burial and physico-chemistry of substitutions, we have designed a predictive model, IPApred (Intrinsically disordered <u>P</u>rotein <u>A</u>ffinity <u>pred</u>iction) (Fig 6). This method allows prediction of the functional effects of single mutations on the binding affinity, in terms of an arbitrary Mutational Tolerance Score (MTS_pred_) for IDPs forming extended interfaces with partners. For each mutant, the MTS_pred_ score is calculated by accounting for the type of substitution, and the residue burial upon binding of the WT residue. The amino acids can be classified into six categories namely charged, polar, aromatic, aliphatic, Proline and Glycine. We have found distinct tolerance patterns for substitutions across these categories in CcdA and calculated category penalties from the CcdA tolerance landscape to account for the effects due to changed chemistry upon mutation. Owing to the fair correlation between positional mutational tolerance and the residue depth upon binding, we have also introduced residue depth to weigh mutational effects at each position. This requires the availability of a complex structure of the interacting proteins. Owing to lack of prolines in the CcdA WT sequence, our model currently does not predict effects of substituting prolines. The IPApred was developed using a training set comprising 60% of the CcdA DMS dataset and was evaluated on a test set of the remaining 40% of the data. The predicted score, MTS_pred_ correlated well (r= 0.65) with the experimental MTS_seq_ for the CcdA mutants test set (Fig 7A). Based on the good correlation between the ΔMTS_pred_ (MTS_pred_^mut^ – MTS_seq_^WT^) values from IPApredand the experimentally determined ΔΔG°_bind_^mut-WT^ in the set of validated twenty CcdA mutants(Fig 7B), we further developed our approach to predict ΔΔG°_bind_^pred^ both for CcdA and other systems. Our predictions for CcdA also correlated moderately with predictions by BeAtMuSiC server (Fig 7C).

**Figure 6.**
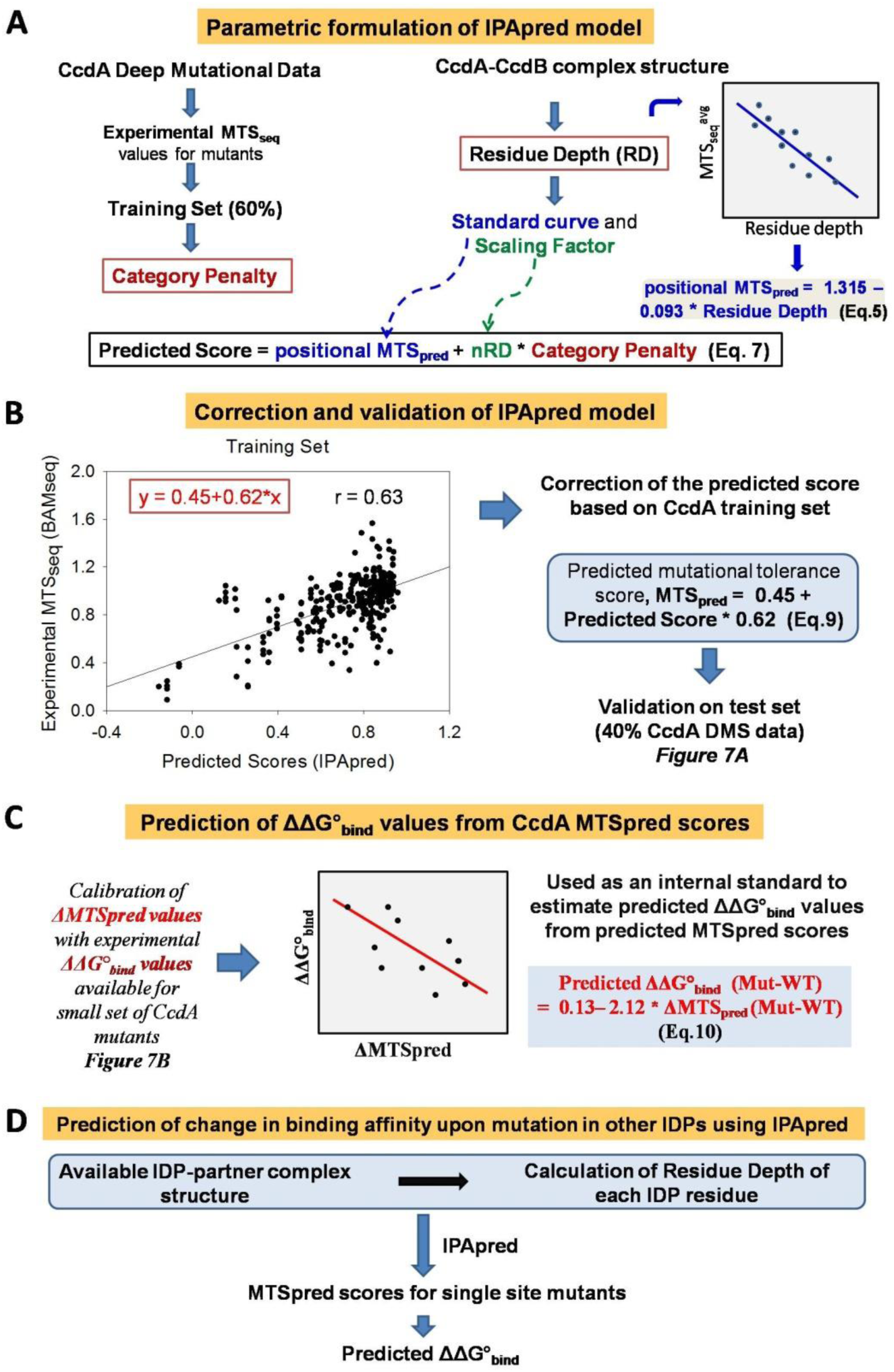
IPApred method development for prediction of binding affinity changes in IDP. (A) The category penalties for each of the 29 categories of substitution, and residue depth (Å) are derived from the CcdA DMS training set and available CcdAB complex structure (PDB id : 3G7Z) respectively. A mathematical formula (equation 7) for estimating relative binding affinities (predicted score) is optimized for best performance using the training set. This uses the positional MTSpred (obtained from dependence of experimental DMS scores on residue depth in CcdA) and category penalty weighted by normalized residue depth. (B) The predicted scores are corrected based on the observed correlation of predicted and available experimental scores in the training set to derive MTSpred which is the final predicted mutational score (equation 9). The method is then validated on the CcdA DMS test set shown in Figure 7A. (C) The linear fit to experimental ΔΔG° binding of twenty individual mutants as a function of ΔMTSpred (MTSpred (mut) – MTSpred(WT)) is used to further estimate the predicted ΔΔG°_bind_ of mutants from MTSpred values (equation 10). (D) Finally the model is applied to other IDP systems. Residue depth is calculated from available IDP-partner structure and category penalties and corrections derived from CcdA are used to predict MTSpred values using the same equations (Eq. 9) as used for CcdA (A-C). The ΔΔG°_bind_ values are then predicted using these calculated MTSpred scores (Eq. 10).

**Figure 7.**
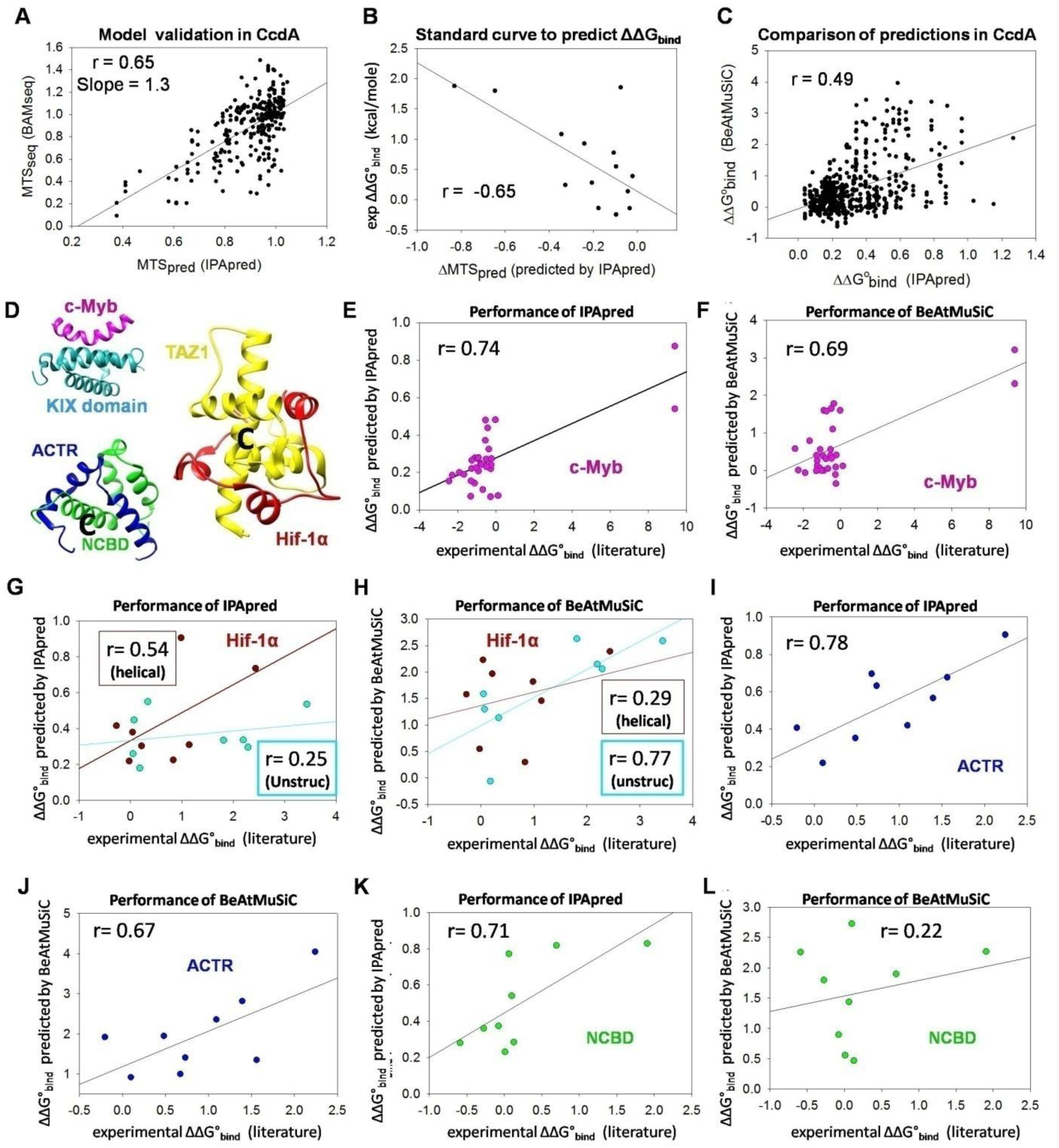
Correlations of experimentally determined mutational effects on binding affinity with predictions by IPApred. (A) Correlation between the mutational tolerance scores predicted by IPApred (MTS_pred_) and the experimental mutational tolerance scores (MTS_seq_) obtained from the DMS study for the test set of CcdA C-terminal domain mutants. (B) Correlation of ΔMTS_pred_ (MTS_pred_^mutant^ - MTS_pred_^WT^) with ΔΔG°_bind_ values determined experimentally for 20 single-site CcdA mutants. The fitted straight line (where y is ΔΔG°_bind_ and x is ΔMTS_pred_) is further used as an internal standard to derive predicted ΔΔG°_bind_ values from MTS_pred_ scores in other systems. (C) Correlation of the ΔΔG°_bind_ values predicted by IPApred for CcdA C-terminal domain mutants in the test set with the corresponding ΔΔG°_bind_ values predicted by BeAtMuSiC for the same mutants. (D) The available structures of intrinsically disordered transactivation domain of c-Myb protein interacting with the KIX domain of CBP protein (PDB id: 1SBO), intrinsically disordered ACTR domain of p160 protein bound to the molten globule like NCBD domain of CBP protein both of which fold into helices upon binding to each other (PDB id: 1KBH), and intrinsically disordered CAD domain of Hif-1α protein interacting with the TAZ domain of CBP protein (PDB id: 1L8C). (E,G,I and K) Correlation of the ΔΔG°_bind_ values predicted by IPApred for disordered protein domains, with available experimentally determined ΔΔG°_bind_ values for corresponding single mutants. (F,H,J and L) Correlation of the ΔΔG°_bind_ values for disordered protein domains predicted by BeAtMuSiC with available experimentally determined ΔΔG°_bind_ values for corresponding single mutants In case of Hif-1α, the mutant dataset was split based on their positions in the helical (brown) or unstructured (blue) stretches in the Hif-1α CAD domain (G-H). The IPApred method appears to perform better for helical stretches than unstructured regions, formed upon TAZ binding in Hif-1α. For ACTR and NCBD, the correlation of predicted and experimental ΔΔG°_bind_ values after removal of a single outlier in each case (indicated in Supplementary Fig S12) is shown (I-L).

Validaion of our affinity prediction model was difficult owing to the lack of adequate and comprehensive literature on single site mutational effects on binding affinity in intrinsically disordered proteins. We therefore proceeded to test the performance of IPApred on the limited datasets of single site mutational effects of small numbers of alanine/glycine/valine substituted variants in disordered protein domains in humans, namely c-Myb(Giri et al., 2013), ACTR (transcriptional co-activator for thyroid hormone and retinoid receptors)(Dogan et al., 2013), NCBD domain of CBP (CREB binding protein)(Dogan et al., 2013) and Hif 1α (hypoxia inducible factor 1α)(Lindström et al., 2018). Our predictions showed better correlation to the literature derived experimental data, relative to predictions of BeAtMuSiC (Fig 7E-L). In case of Hif 1α protein, the disordered C-terminal activation domain forms helical stretches interrupted by large unstructured regions upon binding to its partner, the Taz1 domain of CBP protein (Fig 7D). We noticed that while IPApred accurately predicted mutational effects in the helix forming regions of Hif 1α, our model performed poorly in case of substitutions at residues that constitute the unstructured regions (Fig 7G-H). While BeAtMuSiC excels at predicting binding affinity changes in structured globular proteins, this does not fare as well for PPIs in case of disordered proteins/regions. In case of ACTR, and NCBD experimental values correlated poorly with the predictions made by our method, IPApred (Supplementary Fig S12), as well as the web server, BeAtMuSiC. This is because of a single outlier in each case, namely the ACTR L32A mutation and the NCBD L77A mutations (Supplementary Fig S12). Upon removal of these two mutants, we find that the correlations improve significantly and that the IPApred methodology consistently fares better than the BeAtMuSiC in these small datasets (Fig 7G-H). However, CcdA DMS data driven IPApred isunable to capture stabilizing mutational effects on binding energetics, since CcdA single mutants rarely show better binding to partner relative to WT. We also observed that the quantitative effects of mutations on ΔΔG°_bind_ were heavily dependent on the length of IDP stretch and interface size of the PPI. Therefore, though MTSpred provides an accurate relative binding affinity estimate (relative to WT), the predicted ΔΔG°_bind_values may besmaller or larger than those observed experimentally for IDPs with larger or smaller interfaces respectively. More systematic studies of binding affinity in large variant libraries of various IDP systems are therefore necessary to understand effects of mutations and interface size on IDP binding energetics and to build more accurate prediction tools.

We also used IPApred to test mutational predictions on PPIs for all single-site mutants in three other intrinsically disordered proteins, human oncoprotein p53 and *Mtb* antitoxins, VapB5 and MazEmt9. These IPApred predicted scores were compared to the respective binding affinity predictions by BeAtMuSiC, in the light of absence of exhaustive experimental binding data for all single mutants of IDPs in the literature. Overall, IPApred prediction shows a fair correlation with the BeAtMuSiC predictions for these large number of variants (Supplementary Fig S13), indicating applicability of IPApred for prediction of binding energetic effects for all possible single-site mutants of IDPs.

## Discussion

Deep mutational scanning of a yeast surface displayed library of single-site CcdA variants reveal that mutations in itsN-terminal DNA-binding domain do not affect the CcdB binding activity, consistent with available structural and functional information. The C-terminal intrinsically disordered domain is sensitive to mutations, especially at sites known to be involved in CcdB binding. The mutational effects at the CcdB interacting sites of CcdA are smaller than those commonly observed in globular proteins, likely due to the highly extended interface for the CcdA-CcdB complex that involves about ~32% of all CcdA residues contacting the CcdB dimer. In contrast, active sites of globular proteins typically encompass smaller interface areas and a significantly lower fraction of interacting residues, for example only ~8% of residues in the globular protein CcdB interact with its cellular target bacterial DNA-Gyrase. In globular proteins, interface (active-site) residues show high sensitivity to aliphatic mutants (Tripathi et al., 2016), which is not seen in case of CcdB-interacting positions in the IDP, CcdA C-terminal domain. CcdB-interacting residues of CcdA behave similarly to buried residues in globular proteins, tolerating aliphatic substitutions but not polar or charged mutations (Tripathi et al., 2016). This possibly originates from the largely hydrophobic nature of CcdA-CcdB interactions, which account for nearly 50% of the ΔASA (De Jonge et al., 2009). However, unlike buried sites in globular proteins which are all typically buried deep in the hydrophobic core, interacting residues in IDP interfaces undergo transient burial upon interaction, residing at low residue depth from the complex surface. This expectedly produces a milder energetic effect of single substitutions in disordered CcdA including at the interacting positions. At positions harboring wildtype, aliphatic or aromatic residues in CcdA, charged substitutions are poorly tolerated. At wildtype charged and polar residues, the tolerance pattern generally lacks any apparent amino acid preference. We also observed no prominent effect of the size of substitutions on the CcdB-binding affinity in CcdA.

Despite the high tolerance to mutations in CcdA, our methodology BAMseq is able to detect small changes in binding affinity in a high-throughput manner by integrating YSD coupled FACS methodology to deep sequencing. Previous YSD FACS based method to estimate binding affinity such as Tite-seq, involves experiments at multiple (~11) ligand concentrations(Adams et al., 2016). In contrast, BAMseq involves sorting the library at a single ligand concentration into bins across a binding : expression axis. Carrying out DMS based binding analysis at multiple concentrations may facilitate accurate measurement of binding energetics in globular proteins where single-site mutations can have large effects on binding affinity. In contrast, in cases like CcdA where K_d_ values for single-site mutants vary over a relatively smaller range of values (0.1-7 nM for CcdA mutants) experiments at a single ligand concentration suffice for accurate measurement of binding affinity to target protein. Based on analysis of theoretical curves of binding : expression fluorescence intensity as a function of Gibbs free energy of binding (and dissociation constant) (Supplementary Fig S5), we found that CcdB (ligand) concentrations in the range of 2-10 nM are suitable for the single ligand concentration approach used in BAM_seq_ to parallelly estimate binding affinities of CcdA mutants. A similar approach can be applied to select a single or minimal number of ligand concentrations for K_d_ estimation in other systems.

A mutational tolerance score (MTS_seq_) calculated for each single mutant describes the mutational effects on IDP-target interactions accurately and elucidates sequence–function relationships in IDPs. Further, this scores also allows parallel and high throughput quantification of binding energetics of all protein variants using an internal standard. While wild type CcdA shows a ΔG°_bind_ value of −13.08 ± 0.05 kcal/mole, ΔG°_bind_ for the CcdA single mutants vary over a range of −11 to −14 kcal/mole.In case of mutations with very low surface expression, the Binding:Expression fluorescence parameter fails to represent the actual affinity to target. While some mutations in globular proteins are known to affect surface expression in YSD studies, we observed no such effects on expression in case of IDP CcdA mutants. Unlike the well-studied mutational effects on protein folding and stability in globular proteins, we find no significant mutational effects on stability, proteolysis and/or levels of protein in the IDP, CcdA. This is evident from our exhaustive screening of the expression levels of CcdA mutants relative to WT, on the yeast surface.

The averaged MTS_seq_ score (across all substitutions) for each residue position provides a good estimate of the residue’s contribution to CcdB binding energetics. We developed a method for identification of functionally important residues in IDPs using the deep mutational scan information, which shows high accuracy, sensitivity and specificity. Surprisingly, evolutionary conservation shows poor correlation with CcdA mutational effects, indicating the potential disadvantages in predicting functionally important residues from evolutionary information in case of disordered antitoxins and other similar protein segments. While there is an overall good agreement between experimental mutational tolerance and the residue’s interactions in the complex, there are a few residues that show extensive residue burial in the complex but are surprisingly tolerant to mutations. These are found either close to the N or C-terminus, or at stretches which have a small number of contiguous interacting positions across the primary sequence. Our findings highlight the caveats associated with assuming all interactions identified from the crystal structure are functionally important, especially in case of high affinity interactions involving large interfaces.

Generally, proline residues are found to be abundant in intrinsically disordered proteins (Mateos et al., 2020). The CcdA C-terminus however lacks any wild type proline residue. CcdB binding was observed to decrease upon introduction of proline at most C-terminal domain positions (excepting the 42, 50 and C terminus 70-72 residue positions), including the CcdB non-interacting sites (Fig 3). Proline is known to disrupt helices except when located at the first three N-terminal positions(Bajaj et al., 2007; Kim and Kang, 1999). This is owing to its lack of the amide hydrogen that forms an essential intra-molecular H bond in helices with the i-4 C=O group. The decreased activity of proline mutants indicates that CcdB binding of CcdA, though mediated by an intrinsically disordered domain, may require structuring into α-helical structures upon binding. A similar disruption in IDP function due to proline substitutions throughout the protein length, has also been reported in an amphiphilic helix forming IDP, α-synuclein (Newberry et al., 2020). However, we find poor correlation between helix propensity and observed mutational tolerance in CcdA. This is consistent with previous findings refuting the pre-requisite of an intrinsic helical tendency for optimal partner binding in IDPs(Rogers et al., 2014). Our results suggest that though the intrinsically disordered CcdA C-terminal domain may not require a transiently populated helical structure in the free form to bind CcdB, permanent disruption in the helical structure formed upon binding by introduction of backbone-modulating proline residues may impede CcdB interaction. We also suggest that though the disorder encoded into the primary sequence of an IDP might be essential for functions like prompt bio-removal, promiscuous binding etc., its contribution to partner binding is minimal.

Low mutational tolerance of the CcdB non-interacting Glycine 63 residue of CcdA to all substitutions helps to identify it as an interface shaping residues. This can be rationalized by the positive Φ value of backbone conformation at this position, which allows the helix (40-62) in WT CcdA to break and the 64-72 extended arm to wrap around and bind the CcdB dimer. While BAMseq successfully identifies the previously unknown, functionally important Glycine 63 residue, it fails to recognize it as a non-interacting residue, lacking physical contact with CcdB. We have located several glycine residues with similar positive Φ torsional angle and putative backbone bending roles, widespread in several type II antitoxins (Table 1).

Despite immense advances in protein structural modeling for globular proteins, prediction of partner bound structure and function of IDPs remains a major challenge (Baek et al., 2021; Chandra et al., 2021; Jumper et al., 2021). In the current study we inferred that the CcdA C-terminal region 40-62 forms a helical structure, based on the ~3.6 residue periodicity observed in the tolerance pattern of charged substitutions. While the mutational pattern of the 63-72 residue stretch also shows periodicity, the periodicity of ~4.1 however suggests a predominantly non-helical stretch. Since the mutations in the N-terminal domain of CcdA show no effect on CcdB binding affinity, BAMseq could shed no light on its local structural features (Supplementary Fig S9A). Based on the mutational data, we suggest that both Cysteine scanning and charged residue scanning mutagenesis might be useful in identifying functionally important positions at extended interfaces, and predicting structure attained upon binding in IDPs (Supplementary Figs S8 and S9). Such Single Residue Scanning methods are comparatively cheaper and simpler than full blown saturation mutagenesis studies. Cys mutants produce relatively smaller effects than charged substitutions making the latter more suitable for rapid identification of interface or secondary structure in IDPs. We have recently carried out charged scanning mutagenesis of another antitoxin, MazE6 from *Mycobacterium tuberculosis* to predict functional residues and structural features(Chandra et al., 2021), though in the absence of an experimental structure, the predictions remain to be validated.

Based on the available exhaustive mutational landscape of CcdB binding to the CcdA protein, we developed a prediction method, IPApred to predict mutational effects on partner binding for single mutations in IDPs. IPApred requires as input an IDP-target complex structure to calculate the residue depth of WT residues. We applied IPApred to a number of IDP domains that fold to predominantly α-helical structure upon partner binding and have extended interfaces. We validated our predictions using experimental binding affinities and compared performance of IPApred to that BeAtMuSiC server for a handful of IDPs where structures are known and experimental binding energetics are available for sets of single-mutant variants. We found IPApred to consistently perform better than BeAtMuSiC in case of these IDRs that attain helical structure upon target binding. Since mutational effects on IDP-target binding affinity appear to be dependent on the size of the binding interface, a method like IPApred that provides arbitrary predictive scores (like MTS_pred_) is a useful approach. The MTSpred scores can be converted to ΔΔG° values by using a small experimental dataset of single mutants of the IDP system involved as an internal standard to obtain a mutational prediction on binding energetics for any single-site mutant. Using CcdA experimental affinity values for the same purpose may produce inaccurate quantitative estimates of ΔΔG° values in cases of some IDPs.

IPApred thus provides arbitrary scores depicting relative binding affinity and apparent ΔΔG°_bind_ upon mutation, calculated using a small set of experimental binding affinity measurements for the system of interest. This helps in accounting for the differences in the extent of mutational effects on binding affinity in IDPs with interfaces of various dimensions. Currently IPApred does not correct for the non-uniform mutational tolerance pattern observed in case of CcdA. We believe predictions can be significantly improved by incorporating high throughput mutational data on binding effects in other IDPs, but such data are currently unavailable.

The CcdA mutational study along with investigations of mutational effects in other disordered antitoxin systems (Chandra et al., 2021) indicate that the severity of mutational effects on partner binding is highly dependent of the size of the interaction interface and strength of the binding affinity in the WT IDP. Studies of more IDP-partner interactions will help develop more accurate models to predict the severity of mutational effects on binding affinity. The current results also highlight how mutational effects on IDP function differ both qualitatively and quantitatively from observed and well documented sequence-function relationships in well-folded, globular proteins. Further DMS studies to elucidate sequence-activity relationships in other IDPs will help to decipher the role of protein disorder in PPIs and to develop improved methodology for prediction of mutational effects on binding and function in IDPs. Structure prediction in proteins with few homologs, which is common in IDPs, remains challenging since even recently developed, highly successful deep learning based methods (Baek et al., 2021; Jumper et al., 2021) work best for multiple sequence alignments, containing large numbers of sequences. Our IDP affinity prediction method in combination with deep learning based predictions of protein-protein complex structure (Jumper et al., 2021; Mirdita et al., 2022) can be a powerful tool to decipher the functional effects of mutations in intrinsically disordered functional domains in proteins, where structural and/or functional information is often largely unavailable.

## Materials and Methods

Brief descriptions are provided below, more details are available in the Supplemental Methods Section

### Cloning and CcdA library preparation

Wildtype and single mutants of CcdA were cloned into the YSD plasmid vector, pETcon (Addgene plasmid # 41522), using two or three fragment homologous recombination(Swers et al., 2004) in yeast *S. cerevisiae* EBY100 strain employing a high efficiency LiAc/ssDNA/PEG chemical transformation method (Gietz and Schiestl, 2007). For preparation of the single-site saturation mutagenesis library of CcdA in pETcon vector, we used primers with NNK codons, and the cloning was carried out using homologous recombination in yeast.

### CcdB Purification and Biotinylation

CcdB was purified using an immobilized CcdA peptide affinity matrix(Tripathi et al., 2016). The purified CcdB protein was then biotinylated using EZ-Link Sulfo-NHS-SS Biotin (Thermo-Fisher). 100 μL of 170 μMCcdB was incubated at 4°C for 2 hours in 1X PBS reaction buffer with 2 mM biotin. The non-reacted biotin was then removed by passing the biotinylated sample through a PD SpinTrap G-25 column (Sigma-Aldrich) with a molecular weight cutoff of 5000 Da. ESI-MS of the biotinylated CcdB sample revealed that the majority of CcdB molecules were biotinylated at 1-3 sites. Biotinylated CcdB was quantified by absorbance measurements and the concentration was verified by Tricine-SDS PAGE using known concentrations of lysozyme protein as a standard.

### Yeast Surface Expression and FACS sorting of CcdA Library

CcdA was expressed and displayed on the yeast surface(Chao et al., 2006). Upon induction, the CcdA displayed on yeast cells was labeled with anti-myc primary antibody and biotinylated CcdB (2nM for sorting and varying concentrations between 0.01pM and 200nM for titration experiments) followed by suitable fluorophore tagged secondary antibodies to probe surface expression and CcdB binding of surface displayed CcdA molecules. To estimate binding affinity, the binding:expression signal was used for sorting the library into vertical gates (bins). To additionally quantify the expression levels, the library was also sorted based on expression signals.

Yeast plasmid was purified from each gated population. The CcdA gene was amplified with primers containing gate specific barcodes before pooling and subjecting to deep sequencing at Illumina NovaSeq6000 150 PE Platform (Macrogen, South Korea)

### Determination of Mutational Tolerance Score (MTS_seq_) for each mutant

NGS data for the CcdA mutant library was processed using an in-house developed pipeline (see Supplementary Methods Section). The number of reads in each gate was obtained for each single mutant after data analysis which was then further normalized before the real abundance of mutants across gate is represented. The read numbers were thus converted to frequency distributions of mutants across gates. (see Supplementary Methods for details on normalization).

The MFI^ratio^or the mean binding: expression ratio fluorescence intensity for a mutant was then calculated as follows:

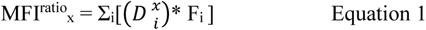

where, x and i represents mutant and gate identity respectively. F_i_ is the mean fluorescence for gate i and 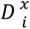 is the normalized fraction of a mutant x in gate i, accurately representing the frequency distribution of mutants across gates (see Supplementary Methods).

TheMFI^ratio^ was calculated for mutants with >100 total reads and averaged across the replicates. The averaged MFI^ratio^ normalized with respect to WT MFI^ratio^ is then assigned as the MTS_seq_(Mutational Tolerance Score derived from deep sequencing) to each mutant. Therefore,

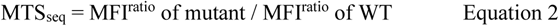

The MTS_seq_ values depicted in the heatmaps are the MTS_seq_ values averaged over replicates 1 and 2 to maximize the information obtainable from the dataset. The MTS_seq_ values used in correlation studies are MTS_seq_ values of Replicate 1, taken from an individual biological experiment and analysis to investigate the performance of single experiment measurements in predicting functional residues etc. The same correlations were also evaluated for MTS_seq_ values of Replicate 2 and showed comparable Pearson correlation coefficients.

Using the same approach, we also calculated the Expression Scores (ES_seq_) for all single mutants of CcdA from the raw mutant reads obtained in the gates P11-P15 (Supplementary Fig S1D).

### Determination of K_d_ and ΔG°_bind_ for surface displayed WT CcdA and single mutants

WT and single mutants of CcdA were cloned in the pETcon vector in *S. cerevisiae* EBY100 strain. The CcdA molecules were expressed, displayed and probed for CcdB binding with a range of concentrations (0.01pM - 200nM) of biotinylated CcdB and Streptavidin conjugated AlexaFluor-633 secondary antibody (1:2000). The mean Alexa-633 (binding) fluorescence was then fitted using SigmaPlot 14, to a one site ligand binding equation :

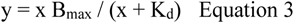

where B_max_ is the saturated Fluorescence Intensity and K_d_ is the dissociation constant of binding (Rathore et al., 2018). The Gibbs free energy of binding (ΔG°_bind_) was calculated using the relationship ΔG°_bind_ = RT lnK_d_, where R (ideal gas constant) = 1.9872 * 10^-3^kcal K^-1^ mole^-1^ and T (temperature) = 298.15 K. The YSD titration estimated K_d_ and ΔG°_bind_ values of CcdB binding for WT and mutant CcdA molecules are denoted as experimentally derived binding energetic parameters (K_d_^exp^ and ΔG°_bind_^exp^) in this work.

### Calculation of apparent ΔG°_bind_ using MTS_seq_ values

Based on the observed high correlation (r = −0.84) between the experimentally determined ΔG°_bind_ values of CcdA molecules (WT and single mutants) and the corresponding deep sequencing derived MTS_seq_ values, we acquired a standard curve describing this relationship. The data points of ΔG° ^exp^versus MTS axis plot were fit to a straight line equation. This was used as an internal standard to calculate the apparent ΔG°_bind_^seq^ from obtained from deep sequencing for mutants (with unknown experimental K_d_ value) using the measured MTS_seq_ values as follows.

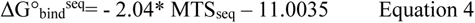

### IPApred method development for prediction of mutant binding affinity changes

For the purpose of designing a predictive model, we used 60% of the CcdA mutational data (comprising of mutant data for the C-terminal domain, residues 40-72) as the training set, and the remaining 40% as the test set to validate the predictive models (Fig 6). The amino acids other than proline and glycine were classified into four categories namely charged, polar, aromatic and aliphatic. Besides these, proline and glycine were considered as individual classes. We calculated category penalties from the CcdA mutational tolerance landscape, to account for the effects due to changed chemistry upon mutation (see Supplementary Methods).We also tested and observed that incorporation of a residue specific penalty (expected to account for residue features like size of the functional side chains)into our predictive model did not improve the predictions. The residue specific penalty was therefore not used in the final model. The good correlation between structure derived residue depth and positional mutational tolerance was used to derive an internal standard curve to estimate the overall sensitivity to mutations for a residue position (positional MTS_pred_) from the residue depth parameter obtained from the protein-partner complex structure.

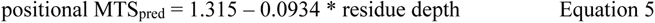

To distinguish between partner interacting and non-interacting residues and to improve the predictor, mutational penalties were weighted by a scaling factor related to the residue depth parameter. After exploring various normalizations of the residue depth, we found the best results using the normalization described below.

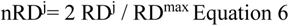

Where RD^j^ is residue depth of a particular residue j in the IDP, and RD^max^ is the maximum residue depth observed in the same IDP segment. The normalization factor of 2 was found to perform best by parametric fitting of equation 6 below for the training set.A number of variations of the model were rigorously tested on the 40% CcdA DMS data (test set). The best performing mathematical formula for prediction of effect of an (a→b) category of mutation at position j, which was finally used in the current work is as follows:

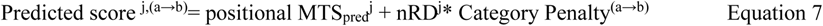

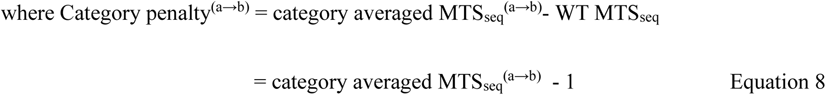

category averaged MTS_seq_ is the averaged MTS_seq_ value for all substitutions of a→b category in the training set.

The predicted score is then corrected based on its observed correlation with the experimental DMS derived MTS_seq_ values for the training set mutants to obtain the predicted mutational tolerance scores, MTSpred (Fig 6).

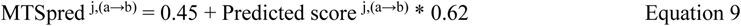

MTSpred values are calculated and validated for single-site mutants in CcdA test set as well as in other IDP systems (Fig 6).

### Prediction of ΔΔG°_bind_ values from predicted Mutational Tolerance Scores

Based on the good agreement between ΔMTS_pred_ (MTS_pred_^mutant^ - MTS_pred_^WT^) andthe experimentally determined ΔΔG°_bind_ values for single-site CcdA mutants, we used this relationship as a standard curve, to derive ΔΔG_pred_ values from the predicted MTS_pred_ values as follows:

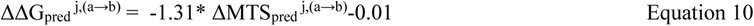

## Author Contributions

R.V and S.C. designed the experiments and wrote the manuscript. K. M. processed the deep sequencing data. A.A helped in acquiring FACS data. S.C. performed all experiments and analyzed the data.

## Acknowledgements

SC acknowledges Ministry of Human Resource Development for her fellowships and thanks all the members of the RV lab for their valuable suggestions. We thank Munmun Bhasin for the useful discussions on protein modelling. We also acknowledge funding for infrastructural support from the following programs of the Government of India: DST FIST, UGC Centre for Advanced study, Ministry of Human Resource Development (MHRD), and the DBT IISc Partnership Program.

## Data and Material availability

The data relevant to the figures in the paper have been made available within the article and in the Supplementary Information section. All materials generated in this study are available from the lead contact without restriction.

