## Supplementary Information for "Mutational scan inferred binding energetics and structure in intrinsically disordered protein CcdA"

#### Supplementary Figures

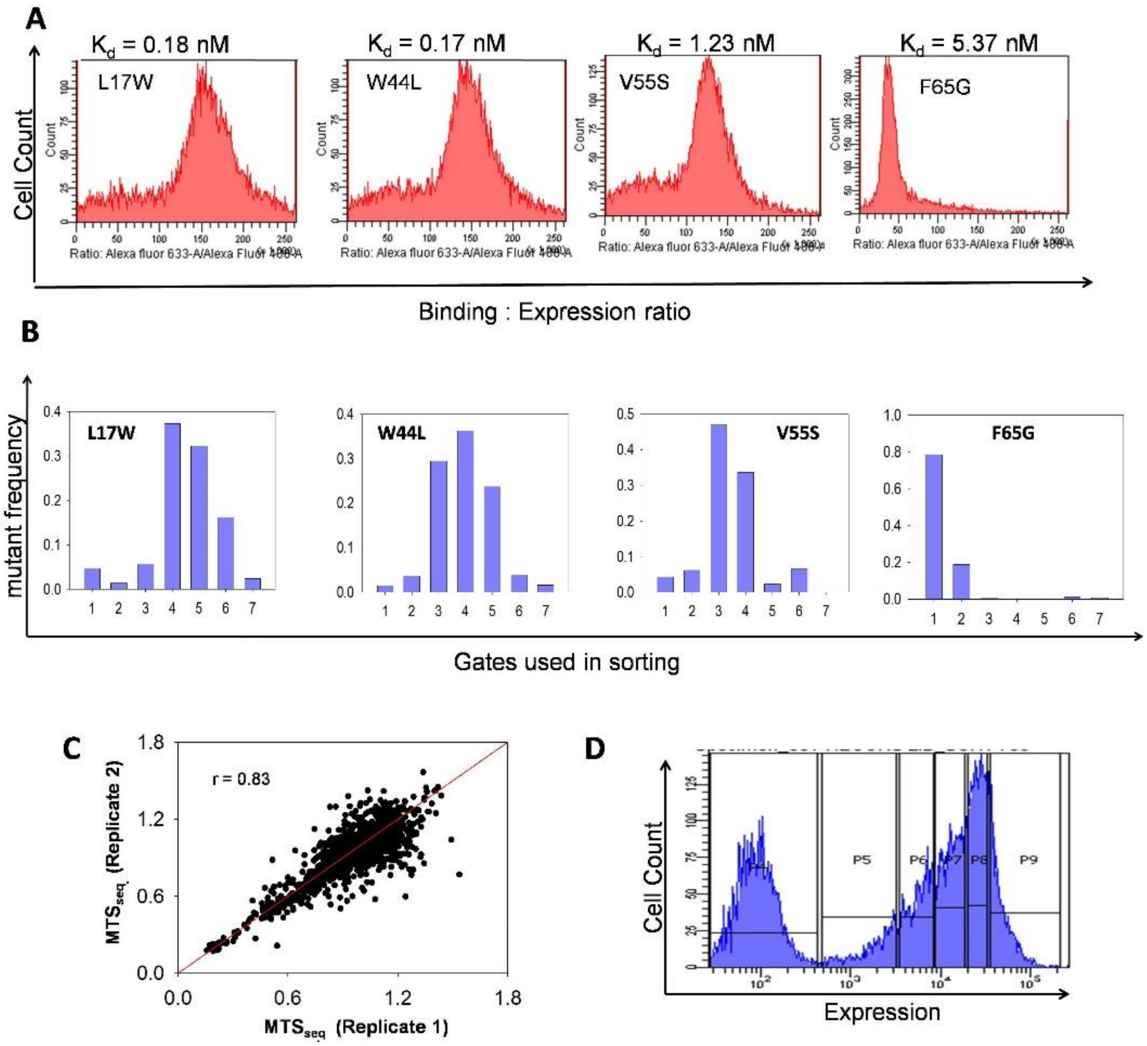


**Supplementary Figure S1. Sorting CcdA library to quantify binding affinity and surface expression** (A) Histograms of Cell count versus Binding : Expression ratio for four single-siteCcdA mutants namely L17W, W44L, V55S and F65G which are known to have distinct CcdB binding affinities listed above (also see Supplementary Table S1). (B) Mutant frequency in each gate used for sorting (in replicate 1) retrieved after analysis of deep sequencing raw read information for the L17W, W44L, V55S and F65G CcdA mutants resembles the count versus binding : expression plot obtained from single experiments (shown in A).(C) Correlation between the MFI$^{ratio}$ values obtained from two biological replicates. The MFI$^{ratio}$ values were calculated for mutants with >100 total reads independently for each replicate. Correlation Coefficient is depicted as r. y=x straight line shown in red. (D) Gates used to sort the CcdA library based on the surface expression (Alexa 488 fluorescence).

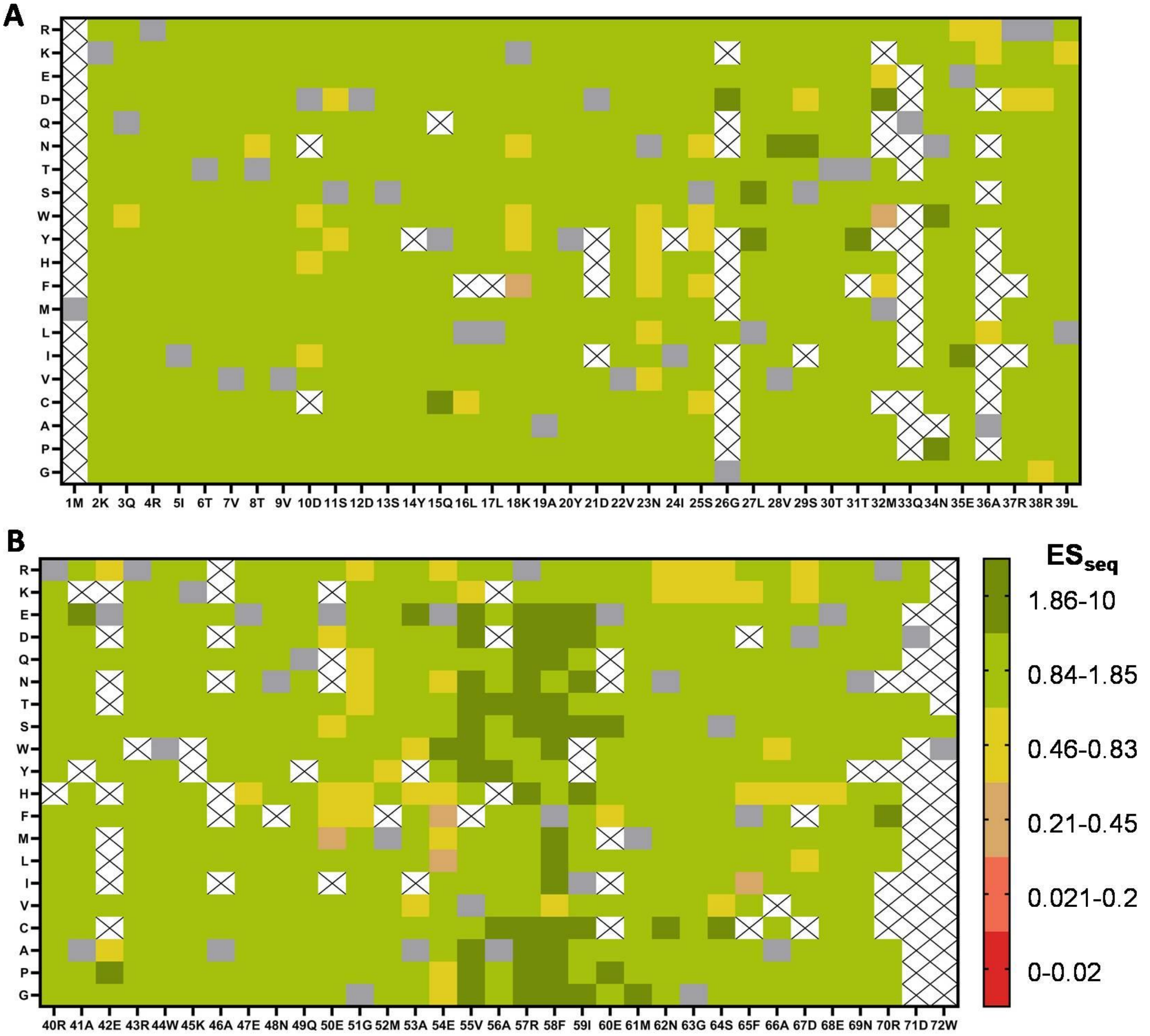


**Supplementary Figure S2. Mutational Effects on Surface Expression for all CcdA mutants** (A) N-terminal domain of CcdA and (B) C-terminal domain of CcdA expression profiles. The x axis depicts the residue number and identity of WT residue, while the y axis refers to the substitutions. Color key depicts the different Expression Scores derived from the deep sequencing, $ES_{seq} = MFI^{exp}$ of mutant / $MFI^{exp}$ of WT. Most single mutants show surface expression comparable to WT ($ES_{seq} = 1$, shown in light green). Positions 55-59 show increased expression. Grey cells and blank cells (marked with X) indicate WT and mutants with no available data respectively.

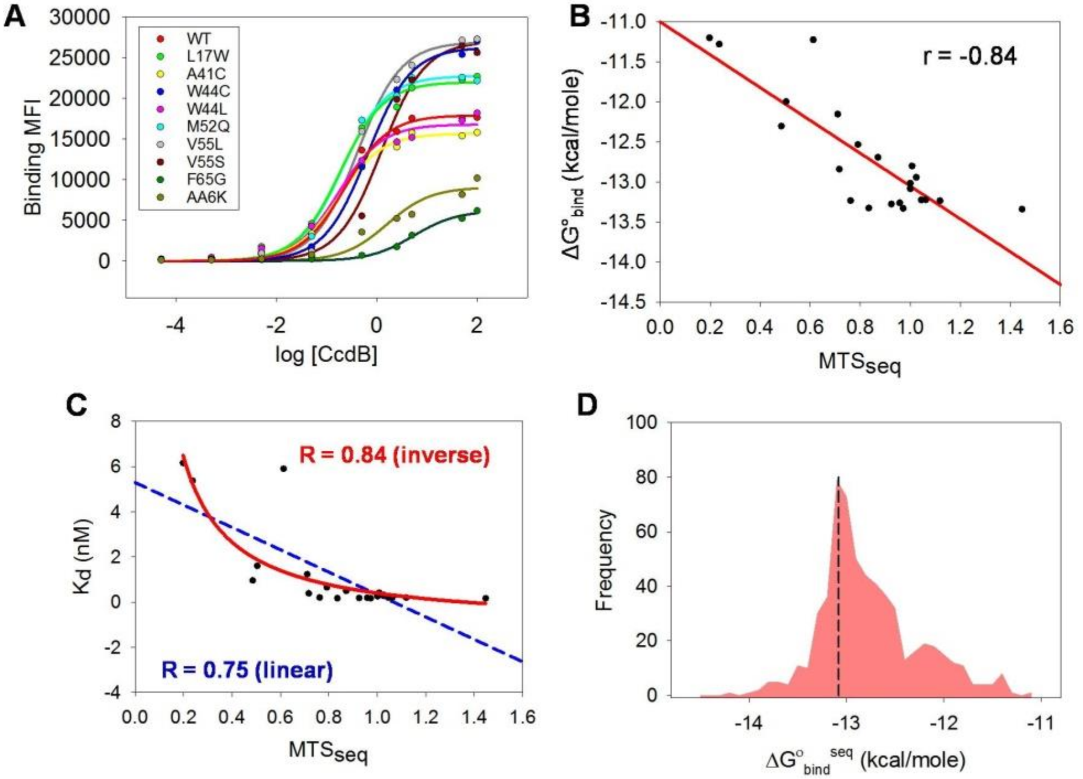


**Supplementary Figure S3. Estimation of ΔG° of binding from Mutational Tolerance Score.** (A) Mean fluorescence intensity (MFI) for CcdB binding as a function of the logarithm of CcdB concentration (in nM) for CcdA WT and nine single mutants using YSD system. Plots for eleven additional single mutants are shown in Supplementary Fig S4. The traces were used to obtain dissociation constant, $K_d$ and $\Delta G°_{bind}$. (B) Correlation between experimentally determined $\Delta G°_{bind}$ for the yeast surface displayed single mutants and the corresponding BAMseq derived mutational tolerance scores ($MTS_{seq}$) respectively. The $\Delta G°_{bind}$ versus $MTS_{seq}$ data was fit to a straight line depicted in red. The fitted parameters were used to estimate the apparent $\Delta G°_{bind}$ (denoted as $\Delta G°_{bind}{}^{seq}$ ) from the $MTS_{seq}$ values for all ~1290 mutants that constitute the CcdA SSM library. (C) The experimentally determined $K_d$ for twenty individual mutants versus the $MTS_{seq}$ scores fitted to a linear equation (blue) and an inverse first order equation, y= c + k/x (red). The good fit for the inverse equation indicates agreement between the theoretical and experimental inverse relationship between the two parameters. (D) Frequency distribution of the $\Delta G°_{bind}{}^{seq}$ for all the CcdA single site mutants estimated as described above. WT CcdA has a $\Delta G°_{bind}$ (experimental) value of -13.08 kcal/mole indicated by a dashed vertical line. The frequency distribution is observed to have a major peak close to the WT value, with a hump at higher range values of $\Delta G°_{bind}{}^{seq}$, comprising of binding defective mutants.

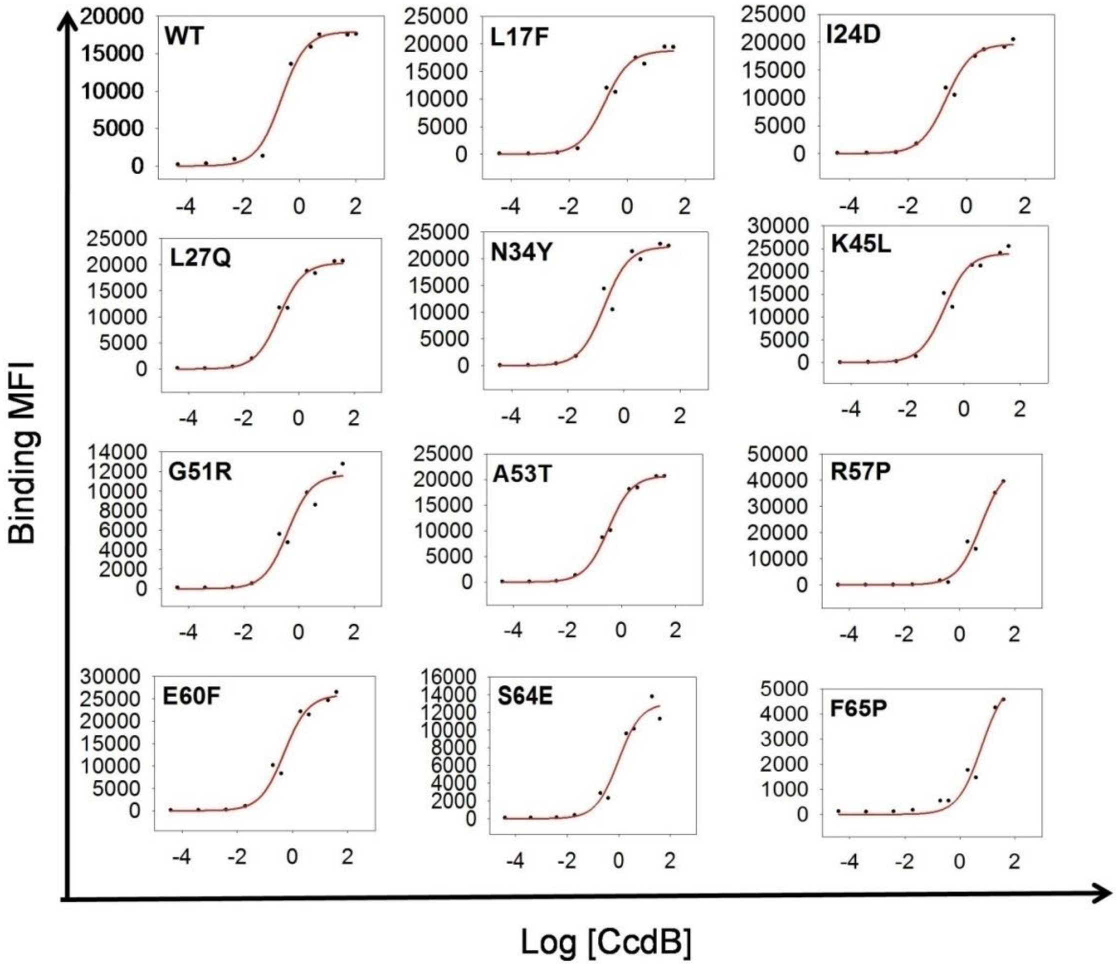


**Supplementary Figure S4.Dissociation constant ($K_d$) measurement for single mutants.** Dissociation constants ($K_d$) were measured for CcdA WT and eleven single-site mutants (other than the nine, shown in Supplementary FigS3A) by titrations followed by FACS analysis of yeast cells displaying CcdA molecules using a range of concentrations (0.01pM-100nM) of biotinylated CcdB. The Mean Fluorescence Intensity values for binding were plotted against the logarithm of CcdB concentration in nM and the traces were fitted to the single-site ligand binding equation , $y = x B_{max} /(x+K_d)$, where $B_{max}$ is the maximal fluorescence intensity, to obtain the $K_d$ (Rathore et al., 2018).

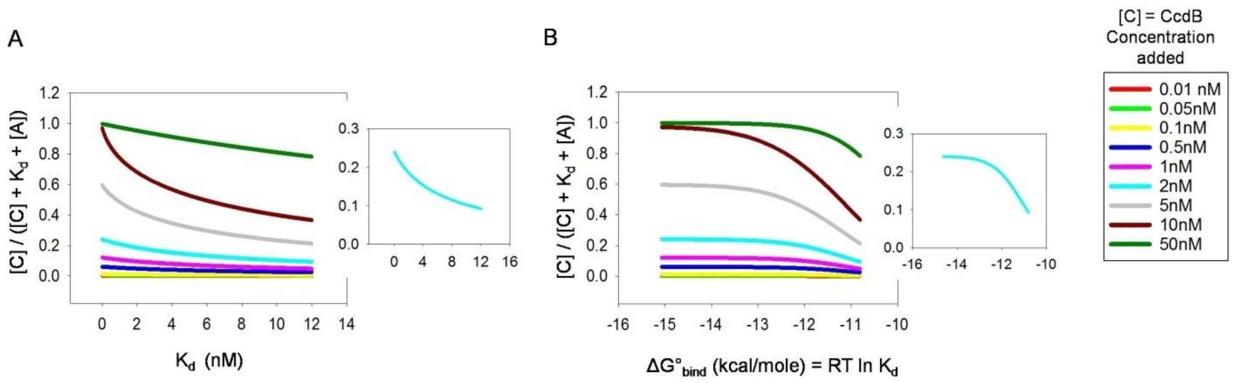


**Supplementary Figure S5. Desirable ligand concentration for BAMseq predicted from theoretical curves.** The MFI$^{ratio}$ is proportional to [C] / ($K_d$ + [C] +[A]) for CcdA YSD (see Supplementary Methods), where $K_d$ = dissociation constant, [C] = total CcdB (ligand) concentration used in FACS experiment and [A] = concentration of unbound CcdA (can be easily calculated using $K_d$ and [C] values). The factor [C] / ($K_d$ + [C] + [A]) is calculated and plotted against a range of $K_d$ values (relevant to the current study) (A) and associated $\Delta G°_{bind}$ values (B), for various ligand ([C]) concentrations depicted in different colors. The theoretical curves at CcdB ligand concentration of 2nM (used in BAMseq) is shown in cyan color and in the insets. These simulated plots indicate that for $K_d$ values in the range 0.1-7 nM (range observed for CcdA mutants), a CcdB (ligand) concentration of 2-10nM is suitable for measuring affinity by BAMseq. At such ligand concentrations, the plots of MFI$^{ratio}$ versus $K_d$ and $\Delta G°_{bind}$ can both be fit empirically to straight lines with significant slopes (though experimental data show better empiric, linear fit in case of $\Delta G°_{bind}$ than for $K_d$ values as depicted in Supplementary Fig S3). This facilitates reasonably accurate estimation of binding affinities for the mutants from the MFI$^{ratio}$ values obtained at a single ligand concentration.

**A**

| | | | | | | | | | | |
|---|---|---|---|---|---|---|---|---|---|---|
| position | 40R | 41A | 42E | 43R | 44W | 45K | 46A | 47E | 48N | 49Q |
| ΔASA | 0 | 38.8 | 0 | 0 | 47.5 | 46.9 | 0 | 0 | 26.5 | 0 |
| $MTS_{seq}^{avg}$ | 0.96 | 0.97 | 0.94 | 0.99 | 0.79 | 0.99 | 1.03 | 1.06 | 1.03 | 1.02 |
| position | 50E | 51G | 52M | 53A | 54E | 55V | 56A | 57R | 58F | 59I |
| ΔASA | 12.8 | 77.1 | 63.6 | 15.4 | 48.4 | 69.3 | 45.6 | 0 | 56.6 | 33.4 |
| $MTS_{seq}^{avg}$ | 0.87 | 0.82 | 0.85 | 0.98 | 0.81 | 0.67 | 0.90 | 1.27 | 0.83 | 0.88 |
| position | 60E | 61M | 62N | 63G | 64S | 65F | 66A | 67D | 68E | 69N |
| ΔASA | 0 | 0 | 20.9 | 2.9 | 25.5 | 96.2 | 85.2 | 18.7 | 31.9 | 62.8 |
| $MTS_{seq}^{avg}$ | 0.85 | 0.99 | 1.11 | 0.83 | 0.77 | 0.39 | 0.68 | 0.77 | 0.83 | 0.81 |
| position | 70R | 71D | 72W | | | | | | | |
| ΔASA | 40.3 | 42.4 | 97.9 | | | | | | | |
| $MTS_{seq}^{avg}$ | 0.91 | 0.89 | 0.94 | | | | | | | |

**B**

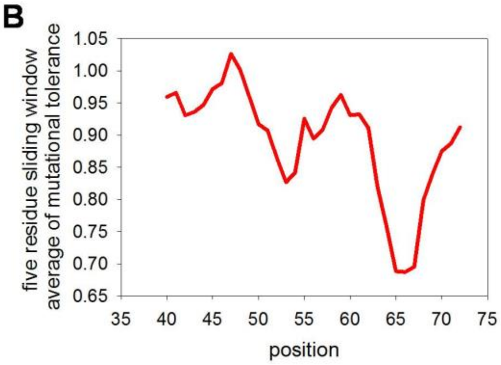


**Supplementary Figure S6. Non-uniform contribution to CcdB binding across the length of CcdA C-terminal domain.** (A) A list of ΔASA (Å$^2$) and the mutational tolerance averaged for all substitutions ($MTS_{seq}^{avg}$), for residue positions in the CcdA C-terminal domain. High ΔASA values highlighted with pink predict structural contacts and low $MTS_{seq}^{avg}$ values highlighted with orange indicate significant contribution to CcdB-binding energetics. (B) Mutational tolerance averaged across a five-residue sliding window and averaged over all 20 substitutions as a function of position. This indicates that contribution to binding is non-uniform across the length of the CcdA C-terminal domain and maximal at residue stretches 52-55 and 64-67.

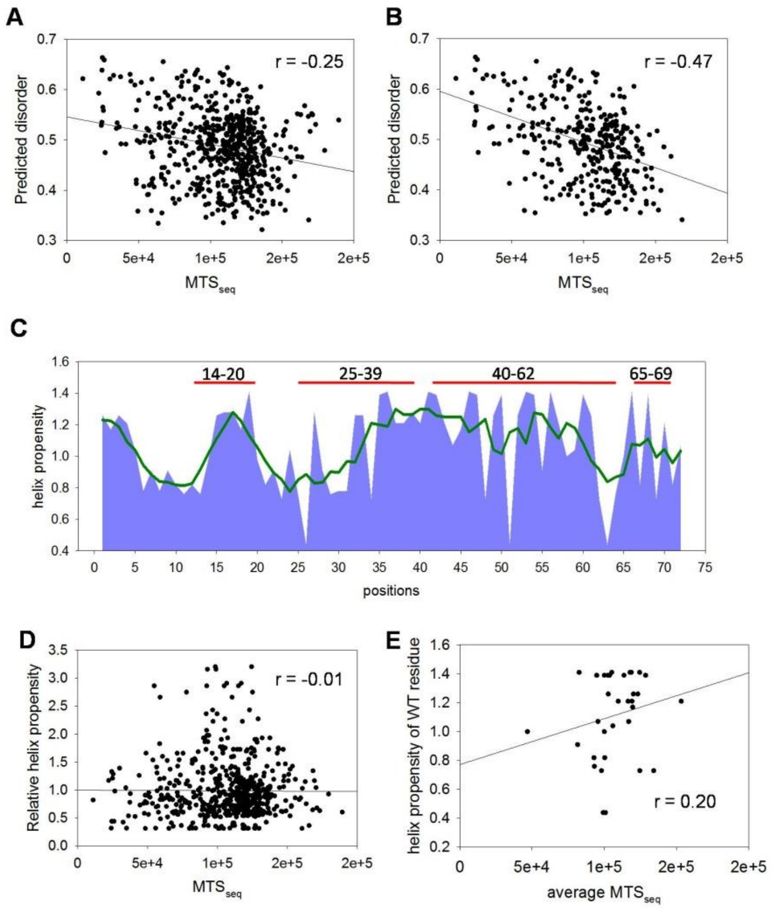


**Supplementary Figure S7. Disorder and helix propensities of CcdA residues are poorly correlated with contribution to CcdB binding** (A) and (B) Correlations between the BAMseq derived $MTS_{seq}$ scores and the disorder predicted by IUPred algorithm (Mészáros et al., 2018) for mutations at all sites (A) and only the CcdB interacting sites (B) in the CcdA C-terminal disordered region. (C) The helix propensity of WT residues across the length of CcdA (blue). The CcdA residue stretches known to form α-helices are highlighted with red lines (De Jonge et al., 2009; Madl et al., 2006). Helix propensity averaged across a five residue sliding window as a function of residue position is shown as olive green line. (D) The relative helix propensity of mutant with respect to the WT amino acid residues for each CcdA single mutant plotted against the respective BAMseq derived $MTS_{seq}$ values. The helix propensity values for the amino acid residues were obtained from the literature (Fujiwara et al., 2012). (E) Correlation between the average mutational tolerance for all substitutions (average $MTS_{seq}$) at each residue position and the corresponding helix propensity of the CcdA wildtype residue.

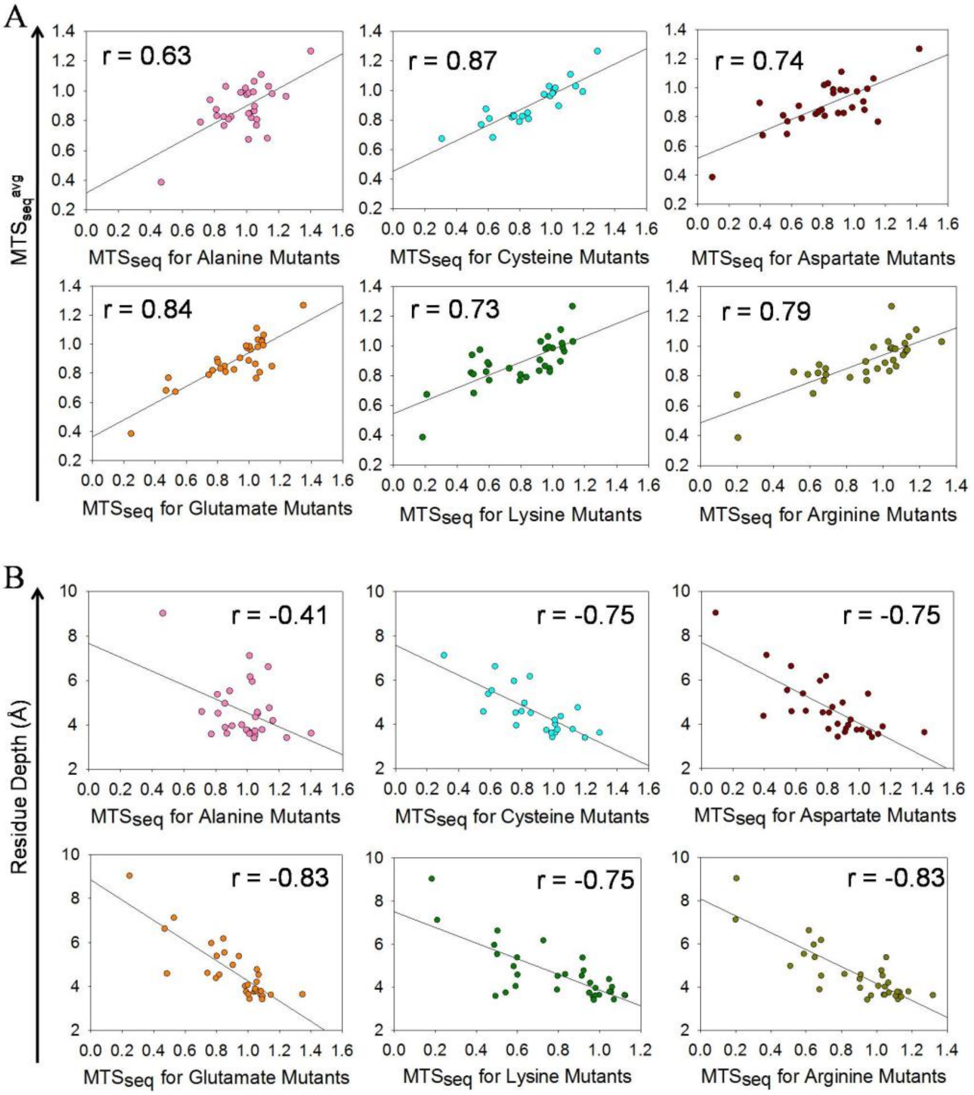


**Supplementary Figure S8. Applicability of Cysteine and Charged scanning mutagenesis to identify interfacial residues.(A)** Correlation of the average mutational tolerance for all substitutions ($MTS_{seq}^{avg}$) at each residue position in CcdA C-terminal domain with the corresponding mutational tolerance of individual Alanine, Cysteine and charged (Aspartate, Glutamate, Lysine and Arginine) substitutions at the same positions respectively. The good agreement between individual residue substitution scores and overall mutational scores suggests the utility of Cys and charged Scanning Mutagenesis methods for interface identification using YSD. (B) Correlation of the residue depth in the CcdAB complex (PDB ID :3G7Z) with the CcdA positional tolerance to Ala, Cys and charged substitutions.Ala mutational effects are found to perform poorly in inferring the functionally important residues in CcdA.

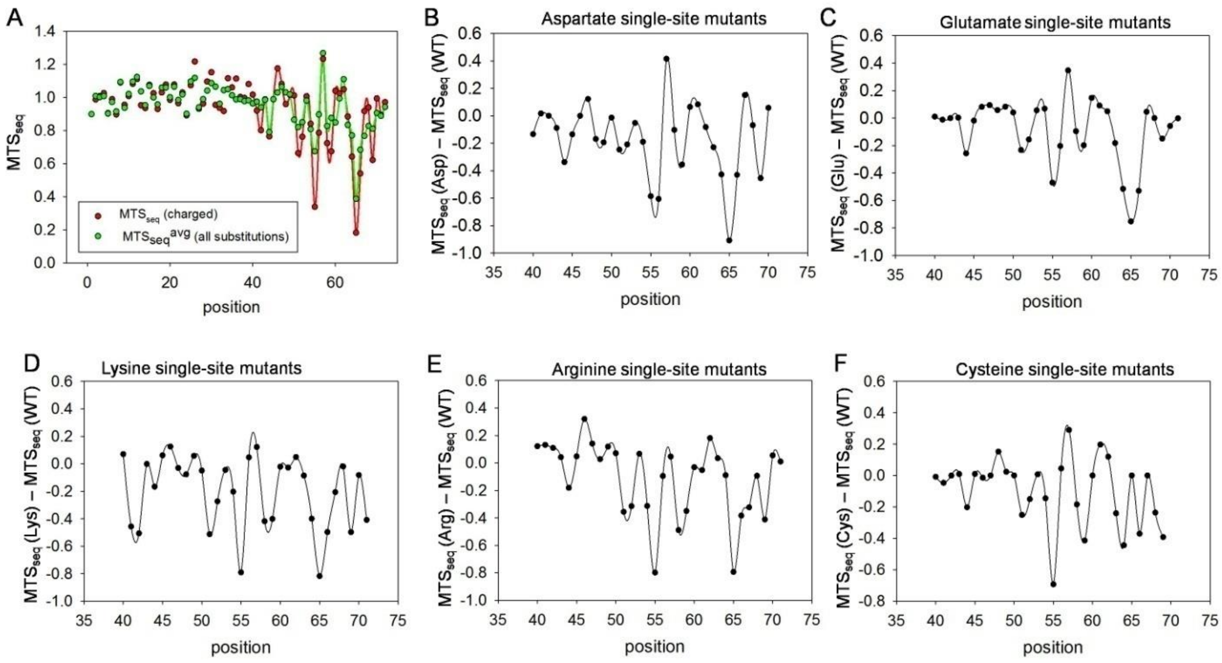


**Supplementary Figure S9. Periodicity in DMS derived mutational scores in C-terminal domain of CcdA.** (A) $MTS_{seq}$ scores averaged over all substitutions (green) and for charged substitutions (red) as a function of all CcdA positions (residues 1-72). Mutational effects are mostly negligible in the N-terminal domain (1-39 residue positions) and thus cannot be used to predict local structural features of the N-terminus. (B-F) Difference between $MTS_{seq}$ scores for single substitutions and WT (where WT $MTS_{seq}$ = 1) as a function of CcdA C-terminal domain positions, shown for all charged substitutions namely Asp, Glu, Lys, Arg and aliphatic, S-containing substitution Cys respectively. The mutational scores for these single substitutions show a clear oscillating pattern that can be used to predict local structural features.

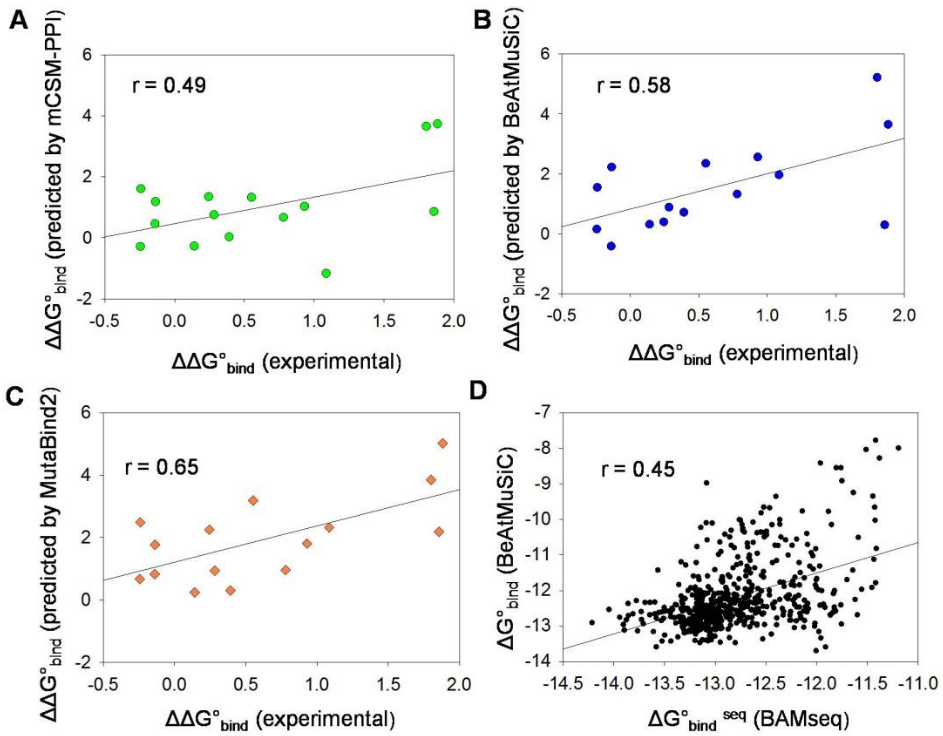


**Supplementary Figure S10. Comparison of experimental CcdA mutational effects on CcdB binding energetics with predictions made by available web servers.** (A-C) Correlations between experimentally determined $\Delta\Delta G°_{bind}^{exp}$ (change in $\Delta G°_{bind}$ upon mutations) values available for the CcdA C-terminal domain mutants and $\Delta\Delta G°_{bind}$ predictions using mCSM-PPI, BeAtMuSiC and Mutabind2 algorithms respectively. Pearson Correlation coefficients are shown in respective plots. (D) Correlation between the apparent free energy of binding, $\Delta G°_{bind}^{seq}$ calculated for all CcdA single mutants from BAMseq and the corresponding $\Delta G°_{bind}$ predicted using BeAtMuSiC server that allows systematic mutagenesis analysis. Most servers considerably overestimate the magnitude of mutational effects on $\Delta\Delta G°_{bind}$ in CcdA.

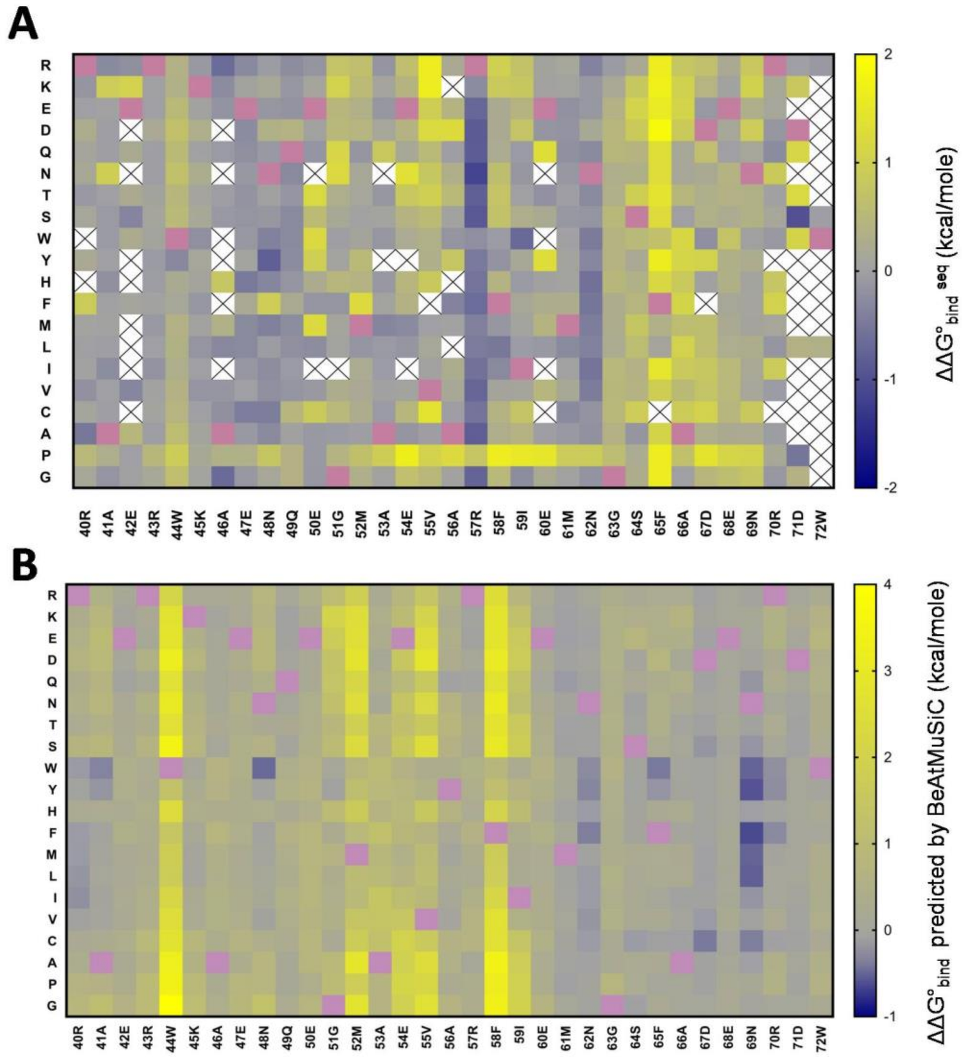


**Sup Figure S11. Comparison of $\Delta\Delta G°_{bind}$ values upon mutations in the C-terminal domain of CcdA, derived from mutational scanning experiments with those from the BeAtMuSiC prediction software** (A) Heatmap of the apparent $\Delta\Delta G°_{bind}^{seq}$ values extrapolated from BAMseq derived $MTS_{seq}$ scores using an internal standard curve. The x axis denotes residue positions in the C-terminal domain of CcdA and the y axis denotes the substituted amino acids. (B) Heatmap of the $\Delta\Delta G°_{bind}$ values predicted by BeAtMuSiC server. Positive $\Delta\Delta G°_{bind}$ (mut-WT) values denote poor binders with lower affinity than WT while negative values denote better binders with higher affinity to partner. Heatmap scales for A and B are non-identical as the predicted mutational effects are more severe than those observed experimentally. Pink cells and blank cells with X denote WT and mutants with inadequate data respectively.

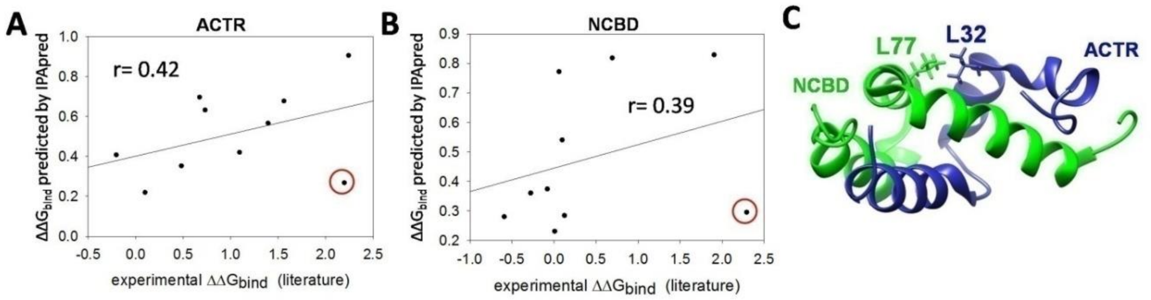


**Supplementary Figure S12. Unusual structural feature of ACTR-NCBD mutants resulting in poor prediction of $\Delta\Delta G°_{bind}$.** Poor correlation of the predicted and experimental $\Delta\Delta G°_{bind}$ values for ACTR and NCBD owing to single outliers in the (A) ACTR and (B) NCBD dataset, (circled in red). These are namely L32A and L77A mutations in the ACTR and NCBD IPDs respectively. (C) The ACTR L32 and NCBD L77 residues mapped on the complex structure of intrinsically disordered ACTR domain of p160 protein bound to the molten globule like NCBD domain of CBP protein. Both unstructured domains fold into helices upon binding to each other (PDB id: 1KBH). These ACTR L32 and NCBD L77 WT residues are found to be fairly exposed, exhibit low residue depth in the complex structure, and are thus expected to be minimally affected by mutation. However, a closer look into the complex structure reveals that despite being surface exposed, these two residues on the respective partners are proximal and possibly involved in interaction with one another, a scenario rising due to the intrinsically disordered nature of both the interacting partners.

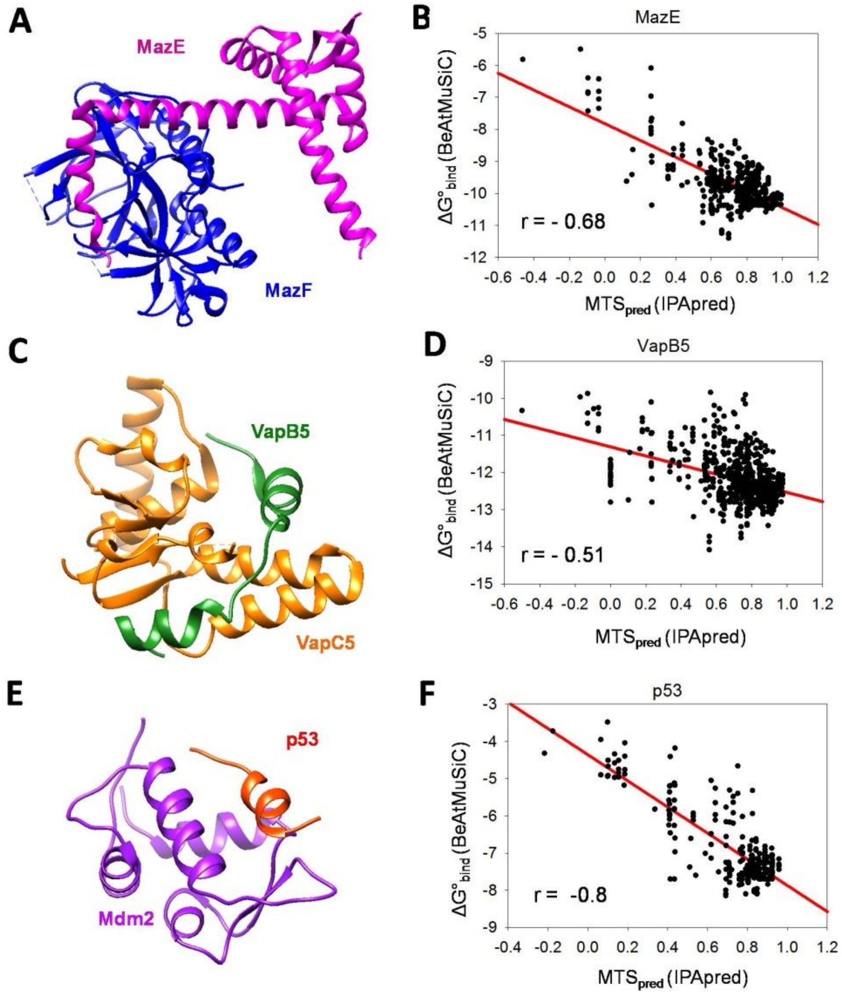


**Supplementary Figure S13.Comparison of predictions of effects on binding affinity upon all possible single-site substitutions in disordered proteins by IPApred with those from BeAtMuSiC software.**(A), (C) and (E) The available structures of partner bound intrinsically disordered domains under study, for the *Mtb* MazEF7(mt9) complex (PDB id : 6A6X), *Mtb*VapBC5 complex (PDB id : 3DB0) and the human p53-Mdm2 complex (PDB id : 1YCR) respectively.(B), (D) and (E) Correlation of $MTS_{pred}$ (arbitrary) scores predicted by IPApred for all possible single mutants, with the predicted $\Delta G°_{bind}$ values from BeAtMuSiC for disordered domains in antitoxins MazE7 and VapB5 and human oncoprotein p53 respectively.

### Supplementary Tables

**Supplementary Table S1. Experimentally determined dissociation constants ($K_d$) and change in Gibb's free energy of binding ($\Delta G°_{bind}$) for WT and single mutants of CcdA using FACS titration**

| Mutant | $K_d$ (nM) | $\Delta G°_{bind}$ (kcal/mole) |
|---|---|---|
| WT | 0.25 ± 0.03 | -13.08 ± 0.1 |
| L17F | 0.16 ± 0.03 | -13.33 ± 0.15 |
| L17W | 0.18 ± 0.02 | -13.27 ± 0.09 |
| I24D | 0.2 ± 0.04 | -13.23 ± 0.16 |
| L27Q | 0.19 ± 0.03 | -13.25 ± 0.13 |
| N34Y | 0.2 ± 0.06 | -13.23 ± 0.25 |
| A41C | 0.17 ± 0.02 | -13.32 ± 0.09 |
| W44C | 0.65 ± 0.05 | -12.53 ± 0.06 |
| W44L | 0.17 ± 0.04 | -13.32 ± 0.2 |
| K45L | 0.2 ± 0.05 | -13.22 ± 0.21 |
| G51R | 0.38 ± 0.12 | -12.83 ± 0.27 |
| M52Q | 0.2 ± 0.03 | -13.22 ± 0.12 |
| A53T | 0.32 ± 0.03 | -12.94 ± 0.07 |
| V55L | 0.41 ± 0.05 | -12.8 ± 0.1 |
| V55S | 1.23 ± 0.19 | -12.15 ± 0.13 |
| R57P | 5.9 ± 1.4 | -11.22 ± 0.2 |
| E60F | 0.49 ±0.1 | -12.69 ± 0.17 |
| S64E | 0.95 ± 0.2 | -12.3 ± 0.17 |
| F65G | 5.37 ± 0.7 | -11.28 ± 0.1 |
| F65P | 6.15 ± 1.7 | -11.2 ± 0.23 |
| A66K | 1.6 ± 0.5 | -11.99 ± 0.27 |

**Supplementary Table S2. Residue burial and Evolutionary Conservation of CcdA C-terminal residues**

| Position and WT amino acid residue | ΔASA (Å$^2$)[a] | Residue Depth (Å)[b] | Percent Evolutionary Conservation[c] |
|---|---|---|---|
| 40R | 0 | 3.5 | 100 |
| 41A | 38.8 | 4.3 | 86 |
| 42E | 0 | 3.4 | 96 |
| 43R | 0 | 3.6 | 97 |
| 44W | 47.5 | 4.9 | 100 |
| 45K | 46.9 | 4.4 | 76 |
| 46A | 0 | 3.7 | 41 |
| 47E | 0 | 3.6 | 97 |
| 48N | 26.5 | 4.7 | 100 |
| 49Q | 0 | 3.8 | 62 |
| 50E | 12.8 | 3.7 | 83 |
| 51G | 77.1 | 6.1 | 100 |
| 52M | 63.6 | 6 | 98 |
| 53A | 15.4 | 4.2 | 74 |
| 54E | 48.4 | 4.4 | 94 |
| 55V | 69.3 | 7.1 | 87 |
| 56A | 45.6 | 4.4 | 100 |
| 57R | 0 | 3.6 | 46 |
| 58F | 56.6 | 5.1 | 100 |
| 59I | 33.4 | 5.3 | 99 |
| 60E | 0 | 3.6 | 95 |
| 61M | 0 | 3.4 | 28 |
| 62N | 20.9 | 3.8 | 97 |
| 63G | 2.9 | 4.5 | 97 |
| 64S | 25.5 | 4.6 | 96 |
| 65F | 96.2 | 8.7 | 100 |
| 66A | 85.2 | 6.9 | 100 |
| 67D | 18.7 | 3.9 | 96 |
| 68E | 31.9 | 3.8 | 83 |
| 69N | 62.8 | 5.5 | 98 |
| 70R | 40.3 | 5.6 | 100 |
| 71D | 42.4 | 4.1 | 66 |
| 72W | 97.9 | 7.5 | 100 |

[a] ΔASA is calculated by subtracting accessible surface area of free structured CcdA by the accessible surface area of CcdA for chain D in complex with CcdB (PDB ID :3G7Z)

[b] Residue depth is a measure of depth of a residue from the bulk solvent in the CcdA-CcdB complex (PDB ID :3G7Z). The values are averaged for CcdA chains C and D in the structure.

[c] Percent Evolutionary Conservation at a position is the percentage occurrence of the *E.coli* wildtype residue in CcdA homologs across all prokaryotes

#### Supplementary Methods

**Safety statement:** No unexpected or unusually high safety hazards were encountered.

**Cloning of CcdA constructs in pETcon vector:** Wildtype and a single site saturation mutagenesis (SSM) library of *ccdA* gene in *E.coli ccdAB* operon were previously cloned in pUC57 vector (Chandra et al., 2022). The antitoxin CcdA gene was amplified from these constructs and cloned into the yeast surface display plasmid vector, pETcon (Addgene plasmid # 41522) using two or three fragment homologous recombination (Swers et al., 2004), in yeast *S. cerevisiae* EBY100 strain using a high efficiency LiAc/ssDNA/PEG chemical transformation method (Gietz and Schiestl, 2007). The pETcon vector was digested with Nde1 and Xho1 prior to recombination. To introduce mutations at the residue positions (40%) which were missing in the starting pUC57 CcdA library, we used three fragment homologous recombination in yeast, where the gene was into split into two PCR fragments with 25 bp homology regions containing the intended mutation. Mutations were introduced using position specific primers containing the NNK codon at the desired position. Single mutants used in titration studies were also individually cloned using three fragments homologous recombination in yeast and were sequence confirmed. Following recombination, plasmid was recovered from yeast, and clones were confirmed by Sanger sequencing of CcdA gene PCR amplified from yeast plasmids of individual clones. The WT pETcon CcdA construct contains a synonymous mutation (GCA→GCG) at A19 residue of CcdA (in comparison to the *E.coli* F plasmid borne *ccdA* sequence) owing to its presence in the starting pUC57 CcdAB construct. The true WT and the A19_GCG synonymous mutant show identical binding affinities to the cognate toxin partner CcdB. The A19_GCG CcdA sequence has been referred to as WT throughout the work.

**CcdB purification :** The CcdB gene, cloned under the pBAD promoter in pBAD24 plasmid and transformed in *E.coli* CSH501 strain (Bernard and Couturier, 1992) was used for CcdB protein expression and purification. 1L LB culture was grown at 37°C and induced with 0.2% (w/v) arabinose at an $OD_{600}$ of 0.5 and grown till an $OD_{600}$of 5. Cells were sonicated at 4°C, after washing and resuspension in HEG buffer (10mM HEPES, 50mM EDTA, 10% Glycerol, pH 7.4). Following centrifugation, the supernatant containing CcdB protein was loaded onto a CcdA peptide (45-72 residues) immobilized Bio-rad Affigel-15 affinity column. After 5 hours incubation at 4°C and washing to remove unbound proteins, CcdB was eluted with 0.2 M Glycine (pH 2.5) into 200 mM HEPES (pH 8.5) at 4°C. After quantification by absorbance measurements using an extinction coefficient, $\varepsilon = 1.4\ mg^{-1}\ cm^{-1}mL$ and confirmation using Tricine-SDS PAGE, the CcdB protein was concentrated to 5 mg/ml and dialysed into 1X PBS (pH 7.4) for storage at -20°C.

**Yeast surface display of CcdA and FACS analysis :** A single colony for CcdA wild–type and single mutants, or 5μL of liquid stock culture for the CcdA Library was inoculated into 5mL of liquid SDCA media and grown at 30˚C and 250 rpm until an $OD_{600}$ of 3-4. Cells were then induced and grown in SGCA media containing 2% galactose at 30˚C and 250 rpm for 16 hrs. After induction, $10^6$ cells were washed with Labeling Buffer containing 1XPBS and 0.5% BSA (kept cold by storing at 4˚C). To monitor the surface expression of CcdA, cells were then incubated for 30 minutes with 20 μL anti-c-myc antibody raised in chicken (1:400 dilution made in Labeling Buffer), that binds to the c-myc tag at the C-terminus of CcdA. The cells were then washed twice and further incubated for 15 minutes with 20 μL secondary antibody Anti-chicken IgG conjugated AlexaFluor-488 (1:300). To monitor the CcdB binding activity of the surface expressed CcdA, the cells were incubated for 30 minutes with 20 μL of 2nM biotinylated CcdB (or varying concentrations between 0.01pM and 200nM for titration experiments). The cells were washed twice following CcdB incubation and incubated for 15 minutes with 20 μL Streptavidin conjugated AlexaFluor-633 secondary antibody (1:2000) that binds to the biotinylated CcdB. The cells were washed thrice after the labeling steps and the now labeled cells were analyzed on a BD FACSARIA III. All incubation steps were carried out at 4˚C at 300 rpm, using a Thermomixer.

Double plots for mean fluorescence intensities of CcdB binding (Alexa-633 Fluorescence) versus Expression (Alexa-488 Fluorescence) were analyzed for the labeled samples. We also used the

Cell Count versus Binding: Expression ratio (Alexa-633: Alexa-488 fluorescence intensity ratio) plots to estimate the relative binding affinities of the different samples.

**Sorting of CcdA Library by FACS:** To examine CcdA mutational effects on CcdB interaction, we analyzed Binding : Expression histograms, where the x-axis (Binding Alexa 633 Fluorescence : Expression Alexa 488 Fluorescence) is a proxy for the binding affinity axis. Cells were sorted into seven vertical gates allowing separation of mutants based on their relative binding activities, in two biological replicates. To assess the mutational effects on surface expression, we also sorted the same library using five vertical gates based on the expression histogram obtained by measuring secondary antibody binding (Alexa 433 fluorescence) to the c-myc tag. The labeled yeast cell library was sorted at the rate of 2,000 events $s^{-1}$ in purity mode and 10,000 cells were collected for each individual sorting gate.

The different sorted populations were collected separately in tubes containing 3mL of liquid SDCA media, and grown to saturation for 36 hours at 30˚C and 250 rpm. Yeast plasmids were extracted from each of the sorted populations after taking equal numbers of cells from each saturated culture. ThermoFischer Scientific GeneJET Plasmid Extraction Kit was used for yeast plasmid extraction, following the manufacturer's protocol with an additional step of overnight incubation in 2U zymolyase at 37˚C before the lysis step. The full length CcdA gene was then PCR amplified from the yeast plasmids, using gate specific barcode containing primers, Phusion Polymerase, and 15 reaction cycles. Following Agarose gel electrophoresis and quantification of DNA concentrations, equal amounts (300 ng) of the PCR products were then pooled and gel band purified using the Gel Extraction Kit. The pooled purified DNA was then subjected to deep sequencing using the Illumina NovaSeq6000 150 PE Platform (Macrogen). Gate specific primers were used to specify the gate identity of the amplified CcdA gene fragments.

**Analysis of NGS Raw data:** An initial data quality check was examined using a PERL script. The paired reads in the raw data were merged using Paired-End Read Merger (PEAR) v0.9.6 tool (Zhang et al., 2014). Merged reads were subjected to "quality filtering", which involves removal of the terminal "NNN" nucleotides, reads with incorrect MID and/or primers, and reads with mismatched MIDs, retaining only those having bases with Phred score ≥ 20. In the binning step, further filtering is carried out to eliminate reads having incorrect primers, truncated MIDs/primers (due to quality filtering) and reads with shorter/longer length than that of the wild type sequences.

Based on the MID, the remaining reads were binned. Each of the binned reads were aligned with the wild type *ccdA* sequence using the Water v6.4.0.0 program(Smith and Waterman, 1981) and reformatted. Finally, after removing reads having indels, they were classified based on the number of substitutions (single, double, triple or multiple).

**Normalization of NGS Raw Reads :** Ideally, the total number of reads in a gate, obtained after deep sequencing data processing should be constant across all gates, since equal numbers of cells are sorted and equal amounts of DNA are processed from each gate. In practice, this does not happen due to experimental errors.

Calculating mutant fractions in each gate helps correct such errors.

Fraction of mutant x in gate i ,

$\left(A_{i}^{x}\right) = \left(a_{i}^{x}\right) / B_i$ Equation S1

where, $\left(a_{i}^{x}\right)$ = number of reads for mutant x in the gate i,

and $B_i$ = total number of reads for all mutants in gate i = $\sum_x \left(a_{i}^{x}\right)$ Equation S2

Since, the initial library population was unevenly distributed amongst the vertical gates, a correction is required, taking into account the population percentage in each gate, to obtain an accurate representation of the mutant frequency distribution.

Therefore, the corrected fraction of mutant x, out of all mutants in a gate i,

$\left(C_{i}^{x}\right) = \left(A_{i}^{x}\right) * P_i$ Equation S3

Where $P_i$ is the percentage of whole library in gate i, as observed in FACS sorting.

We aim to reconstruct a frequency distribution across gates for each mutant, representing the actual abundance of the mutant across the sorted gates in a way that allows easy comparison amongst different mutants. This requires normalizing the data such that all mutants are equally represented. We therefore calculated the normalized fraction of a mutant in a gate assuming identical abundance of each mutant in the whole library population.

Normalized fraction of a mutant x in gate i,

$\left(D_{i}^{x}\right) = \left(C_{i}^{x}\right) / E^x$ Equation S4

where, $E^x = \Sigma_i \left(C_{i}^{x}\right)$ Equation S5

= representation of a mutant x in the whole population, after correction for the uneven population distribution across gates.

The $D_i^x$ terms accurately represent the frequency distribution of mutants across gates, in a fractional form such that for any mutant x ,$\Sigma_i(D_i^x) = 1$. This fractional frequency distribution of a mutant across gates is then used to calculate the mean of the distribution. Since the x-axis, across which gates have been placed, represents Fluorescence Intensity (Binding: Expression ratio), the calculated mean of the distribution is denoted as Mean Fluorescence Intensity (MFI).

$$MFI^{ratio}_x = \Sigma_i[(D_i^x) * F_i] \qquad \text{Equation 1}$$

where, x and i represents mutant and gate identity respectively. $F_i$ is the mean fluorescence for gate i recorded during FACS experiments.

**Calculation of Accessibility, Depth, CcdB Interactions and Evolutionary conservation :**

The accessible surface area, number of interactions with CcdB and residue depth was calculated for each residue in the C-terminal domain of CcdA (C chain in pdb file) using the CcdA-CcdB complex structure (PDB: 3G7Z)(De Jonge et al., 2009). The accessible surface area (ASA) was calculated using the in-house psa program with a 1.4 Å probe radius(Chakravarty and Varadarajan, 1999; Lee and Richards, 1971). The ΔASA values were calculated for each C-terminal residue in chain D in CcdAB structure (PDB id: 3G7Z) using the formula:

$$\Delta ASA^{residue} = ASA^{residue}\text{ (free structured CcdA)} - ASA^{residue}\text{ (CcdB-bound CcdA)} \qquad \text{Equation S6}$$

The residue depth was calculated using the DEPTH server (Chakravarty and Varadarajan, 1999; Tan et al., 2011, 2013). For each residue the residue averaged depth of all atoms was considered. Averaged values for CcdA chains C and D in CcdAB complex structure (PDB id :3G7Z) were used for the residue depth values in the current study. The detailed interaction information including the number of hydrogen bonds and non-bonded interactions each residue forms with CcdB, was obtained from PDBsum(Laskowski et al., 2018).

To calculate evolutionary conservation in CcdA, all proteins annotated as bacterial antitoxin CcdA were extracted from the NCBI gene database. The resulting fasta sequences of CcdA were then aligned using ClustalX and this multiple sequence alignment was fed to the ConSurf server to obtain the residue conservation scores(Ashkenazy et al., 2010). The percent conservation of the *E.*

*coli* CcdA WT residues have been referred to as evolutionary conservation scores in the current work.

A list of ΔASA, residue depth and evolutionary conservation thus calculated for each CcdA residue is available in Sup Table S2.

**<u>Estimating $\Delta\Delta G°_{bind}$ using available computational prediction methods :</u>**To compare our $\Delta G°_{bind}$prediction for mutants with available methods, we used three computational methods to predict the CcdA mutational effects on CcdB binding energetics. Predicted values of $\Delta\Delta G°_{bind}$ ($\Delta G°_{bind}{}^{Mut}$- $\Delta G°_{bind}{}^{WT}$) were obtained from available servers BeAtMuSiC(Dehouck et al., 2013), mCSM-PPi(Rodrigues et al., 2019) and MutaBind(Li et al., 2016).

BeAtMuSiC was also used to predict $\Delta\Delta G°_{bind}$ for all single mutants in the interacting domains of CcdA (residues 40-72), p53 (residues17-29), MazEmt9 (residues56-76) and VapB5 (residues53-86). The PDB IDs for the structures used in the above systematic mutation effect prediction by BeAtMuSiC were 3HPW(De Jonge et al., 2009) , 1YCR(Kussie et al., 1996), 6A6X(Chen et al., 2019) and 3DBO(Miallau et al., 2009) respectively.

BeAtMuSiC was also used to predict $\Delta\Delta G°_{bind}$ for the single mutants in the disordered protein domains of ACTR, NCBD, c-Myb and Hif-1α.

**<u>IPApred model development</u>**

1. <u>Calculation of mutational penalties:</u> To account for mutational effects due to altered physico-chemical properties of substitutions, we calculated penalties specific to the different types of substitutions and used these penalties to predict the mutational effects of substitutions. Amino acids were classified into six classes namely proline (P), glycine (G), aliphatic (A,C,V,I,L,M), aromatic (F,H,Y,W), polar (S,T,N,Q) and charged (D,E,K,R)(Tripathi et al., 2016). Therefore, mutations (WT to Mut) could be classified into 36 categories (6 X 6 = 36), of which Pro→Pro and Gly→Gly can be disregarded, making a total of 34 possible categories of substitutions based on our amino acid classifications. Since CcdA has no WT proline residues, we could not extend our model to the five categories describing substituting proline residues to the other classes. $MTS_{seq}$ values for substitutions falling under each of the remaining 29 categories were averaged to obtain

average category $MTS_{seq}$. The $MTS_{seq}$ values used here were derived from the training set comprising of 60% CcdA DMS results. Category penalties for each of the 29 categories in CcdA were thus calculated by subtracting the category averaged $MTS_{seq}$ from the WT $MTS_{seq}$ as follows:

Category penalty = category averaged $MTS_{seq}$ - WT $MTS_{seq}$

= category averaged $MTS_{seq}$ - 1 Equation 8

Similarly, residue specific penalties were calculated that help describe the amino acid specific mutational effects within the same class, if any.

Residue penalty (of amino acid i)= averaged $MTS_{seq}$ for substitution i across all positions - WT $MTS_{seq}$ = averaged $MTS_{seq}$ for substitution i across all positions –1 Equation S7,

The averaged $MTS_{seq}$ for substitution i = $(\sum_j MTS_{seq}{}^{i})/ N$ Equation S8,

Where i refers to each of the 20 substitutions and N refers to the number of residues in CcdA C-terminal domain with available $MTS_{seq}$ scores for a substitution i.

Using the above equation, we calculated residue penalties for all the twenty amino acids from the CcdA DMS data. The CcdA DMS dataset (for the intrinsically disordered and CcdB interacting C-terminal domain residues, 40-72) was randomly divided into two subsets, the training set and the test set comprising of 60% and 40% of all mutants respectively. The penalties described above were calculated from the training set.

2. Standard Curve for estimating mutational sensitivity from residue burial information:

CcdB-interacting positions in CcdA with high residue burial in the complex were observed to be more sensitive to mutations than the non-interacting sites in CcdA which have low residue burial. Residue depth describes the burial of each CcdA residue in the CcdAB complex (PDB id : 3G7Z). The residue depth correlates well with the average mutational tolerance for CcdA. The fitted straight line equation is used as a standard to estimate the average predicted MTS for each position (positional $MTS_{pred}$), which describes an overall position specific tolerance to mutations.

Positional $MTS_{pred}$ = 1.315 – 0.0934 * residue depth Equation 5

3. Scaling and correcting the category penalties: To predict the mutational effect on partner binding, the category penalties are then weighted by the normalized residue burial, since we find the extent of mutational effects to be dependent on the contribution of the WT residue to binding.

Corrected Category Penalty = Category penalty * $nRD^j$ Equation S9,

Where $nRD^j = 2\ RD^j / RD^{max}$ =normalized residue depth (Å) of position j Equation 6,

And $RD^j$ = residue depth (Å) of WT residue at position j obtained from the available complex structure and $RD^{max}$ is the maximum residue depth observed in the same IDP segment.

The normalized residue depth is a fraction that allows weighing the category penalties based on the extent of interactions the position makes with the partner.

The predicted score is then calculated by adding the corrected category penalty to the positional $MTS_{pred}$.

Predicted score= positional $MTS_{pred}$ + Corrected Category Penalty

= positional $MTS_{pred}$ + $nRD^j$* Category Penalty Equation 7

The category penalties were calculated on a training set using 60% of the CcdA Deep sequencing mutational data and evaluated on a test set comprising the remaining 40% of the data. This predictive approach is currently unable to predict effects of mutations at positions harboring wildtype proline residues, since CcdA lacks any wild type proline.

The predicted scores show good correlation with the BAMseq derived mutational scores, $MTS_{seq}$ for the training set (r = 0.63) (Fig 6). Using the fitted line as an internal standard, we correct the predicted scores to obtain final score, $MTS_{pred}$ (Fig 6).

$MTS_{pred}$ = 0.45+0.62*predicted score Equation 9

**Calculation of Residue Depth:** The residue depth of all positions in the query protein (IDP domain) was calculated using the listed complex PDB IDs as inputs from the available DEPTH web server (Chakravarty and Varadarajan, 1999; Tan et al., 2011, 2013). The residue depth (Å) averaged over all atoms for each residue position in the complex structure was used as input for the predictive model, IPApred.

**Prediction using IPApred:** The residue positions (all or positions with experimental information, depending on the kind of validations) in an intrinsically disordered domain that is involved in partner binding, were used for residue depth calculations. Residue depth for each of these residue positions was obtained as described previously, and these values were input into the mathematical formula used in the IPApred model, Equations 9. For each mutation, the category of substitution (based on the class of WT and mutant amino acid residues) was determined and the respective penalties were calculated from CcdA mutational data for the C-terminal domain. The resulting $MTS_{pred}$ scores, were then used to estimate the predicted values of free energy changes upon binding ($\Delta\Delta G°_{bind}{}^{pred}$). To obtain the $\Delta MTS_{pred}$ for mutants in case of any IDP system, we subtracted the mutant $MTS_{pred}$ by the value of WT $MTS_{seq}$ value (=1) for CcdA data.

**List of PDB IDs used in the study:** The available structures of CcdA - CcdB complex (PDB id: 3G7Z)(De Jonge et al., 2009), c-Myb - CBP complex (PDB id: 1SB0)(Zor et al., 2004), ACTR – NCBD complex (PDB id: 1KBH)(Demarest et al., 2002), Hif-1α – CBP complex (PDB id: 1L8C)(Dames et al., 2002), p53 – Mdm2 complex (PDB id: 1YCR)(Kussie et al., 1996), *Mtb*Maz EF (mt9) complex (PDB id: 6A6X)(Chen et al., 2019) and *Mtb*Vap BC 5 complex (PDB id: 3DBO)(Miallau et al., 2009) were used in this study.

**Derivation of relationship between experimental fluorescence intensity and dissociation constant, $K_d$**

CcdA (A) and CcdB (B) binding and formation of the CcdA-CcdB complex (AB) on the yeast surface can be represented as:

$A+B \leftrightarrow AB$

The dissociation constant, $K_d = [A][B]/[AB]$ Equation S10

The binding fluorescence signal (MFI of binding) is proportional to the number of CcdB-bound CcdA molecules

MFI of binding $\propto$ [AB] Equation S11

The expression fluorescence signal (MFI of expression) is proportional to the total number of CcdA molecules on the yeast surface

MFI of expression $\propto [A]_T$ Equation S12

The ratio of binding : expression fluorescence signal ($MFI^{ratio}$) is proportional to the fraction of bound CcdA

$MFI^{ratio} \propto [AB]/[A]_T$

Or, $MFI^{ratio} \propto [AB]/[AB]+[A]$ Equation S13,

Combining equations S10 and S13, we get

$MFI^{ratio} \propto [B]/ (K_d + [B])$ Equation S14,

Therefore, our calculated binding: expression ratio scores are inversely proportional to dissociation constant, $K_d$ values. Therefore, a linear regression fit is not suitable for the relationship between MTS values and $K_d$ values. On the other hand, when the $K_d$ values are plotted in logarithmic scales we find an empirical linear relationship with the ratio scores. Thus we proceeded to use the free energy of binding ($\Delta G°_{bind} = RT \ln K_d$ ) to obtain a standard curve between experimental binding constants and the mutational tolerance scores (MTS).

The binding: expression ratio score is also dependent on free CcdB concentration in the solution [B]. [B] in turn is related to $K_d$ and total concentration of CcdB (ligand) [C] added, where [C] = [B]+ [AB] Equation S15

Following equation S10 and equation S15, we get,

$[B] = [C] K_d / ([A] + K_d)$ Equation S16

Combining equation S14 and equation S16, we therefore get,

$MFI^{ratio} \propto [C] / (K_d + [C] + [A])$ Equation S17

The [A] values can be calculated using equation S10 and total concentration of CcdA ($[A]_T$). Since, $10^6$ yeast cells that are expected to display approximately $10^5$CcdA molecules per cell(Boder and Wittrup, 2000), are used in a 20 μL reaction for FACS experiments, we assume that the total CcdA concentration, $[A]_T$= 8.3 nM.

When ([C] / ($K_d$ + [C] + [A]) values are simulated for ranges of ligand concentrations, [C]and plotted against $K_d$ or $\Delta G°_{bind}$ values (Sup Fig S5), we can predict suitable CcdB concentrations [C] to be used in experiments for estimation of $\Delta G°_{bind}$ values for large libraries of variants.
